# Prion-like domains control plastid targeting of PEPSI effectors and PEPSI6-mediated modulation of DXR during root symbiosis

**DOI:** 10.64898/2026.05.15.725075

**Authors:** Ernesto Llamas, Thomas Kell, Maurice Koenig, Cecile Angermann, Seda Koyuncu, Jana Kastner, Dennis Mahr, Parisa Kakanj, Taim Nassr, Gerd U. Balcke, David Vilchez, Gregor Langen, Björn Heinemann, Tatjana M. Hildebrandt, Johana C. Misas Villamil, Alga Zuccaro

## Abstract

Microbial effectors often act in the apoplast or cytosol, but how they reach host organelles during beneficial plant–fungal interactions remains poorly understood. Here, we identify a class of secreted prion-like effector proteins from the mutualistic root endophyte *Serendipita indica*, termed PEPSIs. These proteins contain prion-like domains (PrLDs) embedded within intrinsically disordered regions, and three tested PEPSIs require these domains for plastid localization. Focusing on PEPSI6, we show that plastid targeting depends on its PrLD and is associated with engagement of plastid import and proteostasis machinery. In plastids, PEPSI6 associates with 1-deoxy-D-xylulose 5-phosphate reductoisomerase (DXR), promotes DXR accumulation, alters methylerythritol phosphate pathway metabolites, and enhances tolerance to the DXR inhibitor fosmidomycin. PEPSI6 also undergoes apoplastic C-terminal CAP-domain processing, and its colonization-promoting activity persists after PrLD deletion, indicating a plastid-targeting-independent function. Overall, this work identifies PrLDs as noncanonical plastid-targeting elements in PEPSI effectors and reveals DXR as a target of fungal manipulation during root symbiosis.

**Highlights:**

- Prion-like domains control plastid targeting of selected PEPSI effectors.
- PEPSI6 associates with DXR and promotes its accumulation in plastids.
- PEPSI6 alters MEP-pathway metabolites and increases tolerance to fosmidomycin.
- Apoplastic processing reveals a plastid-independent PEPSI6 activity during root symbiosis.

## Introduction

Plant-associated fungi deploy diverse effector proteins to manipulate host cellular processes and promote colonization. While many effectors act in the apoplast or cytosol ^1,2^, an emerging topic in both plant and animal systems is that microbial effectors and virulence factors can also target host organelles to modulate immunity, metabolism, and signaling ^3^. In plants, plastids function as central metabolic and immune hubs that coordinate photosynthesis, redox homeostasis, nucleotide metabolism, hormone biosynthesis, and defense-associated signaling ^4^. This integrative role makes plastids attractive, yet still largely underexplored, targets for microbial interference. Recent work indicates that effectors from pathogenic fungi and beneficial endophytes can alter plastid protein import, stability, and signaling to reshape host immunity. In *Serendipita indica*, for example, the effector *Si*E141 binds a plastid-targeted oxidoreductase and prevents its import, thereby rewiring salicylic acid–dependent defense responses ^5^. Evidence from arbuscular mycorrhizal (AM) symbioses further suggests that plastid-associated defense and metabolic pathways are reprogrammed during colonization ^6^, raising the possibility that plastids represent strategically important organelles targeted across mutualistic interactions. Despite these insights, the mechanisms by which fungal effectors access plastids and directly modulate plastidial metabolism remain poorly understood. In particular, few generalizable signals or domains that enable microbial effector trafficking into host organelles have been identified, and direct effector-mediated modulation of plastid-resident metabolic enzymes remains mechanistically unexplored. Addressing this gap is essential for understanding how mutualistic fungi reprogram host metabolism and immunity at the organelle level.

*S. indica* is a root-colonizing endophyte capable of establishing beneficial associations with a wide range of plant species, including *Arabidopsis thaliana* ^7–10^. Its colonization strategy involves an initial biotrophic phase followed by a cell death–associated phase, supported by numerous secreted effectors that modulate oxidative stress, host cell death, metal homeostasis, and immune responses ^5,11–16^. Although hundreds of candidate effectors are predicted in the *S. indica* genome ^17,18^, only a small subset has been functionally characterized, and the mechanisms by which intracellular effectors engage plastids and other host organelles remain largely unknown.

One potential clue lies in the structural features of the effectors themselves. Recent work has shown that plant cells can recognize and process proteins with prion-like or highly disordered domains through the chloroplast import and proteostasis pathways ^19^. These findings raise the possibility that unconventional structural elements within fungal effectors might similarly facilitate organelle targeting. Intrinsically disordered regions (IDRs) are protein segments that lack stable tertiary structure yet remain highly dynamic and functional. Their conformational plasticity enables interactions with multiple partners, responsiveness to cellular conditions, and the assembly of biomolecular condensates ^20,21^. A subset of IDRs contains prion-like domains (PrLDs), enriched in uncharged polar residues, particularly glutamine and asparagine, that drive liquid–liquid phase separation, formation of higher-order complexes, and unconventional trafficking behaviors. Although PrLDs have been studied extensively in RNA-binding proteins and stress-responsive regulators across eukaryotes, their roles in fungal effector biology remain unexplored. Through genome-wide analyses, we identified a group of secreted *S. indica* proteins that contain PrLDs and are upregulated during root colonization, which we refer to as <u>p</u>rion-like <u>e</u>ffector <u>p</u>roteins of *<u>S</u>. indica* (PEPSIs).

Here, we show that three PEPSI proteins localize to host plastids and that their central PrLDs, embedded within IDRs, are required for plastid localization. To investigate the underlying mechanism, we characterized PEPSI6, which contains an N-terminal IDR harboring a central PrLD and a C-terminal predicted CAP domain. CAP (Cysteine-rich secretory protein, Antigen 5, and Pathogenesis-related 1) domains are conserved structural modules found across eukaryotes and, in plants, are best known for generating CAPE peptides, bioactive signaling molecules released in the apoplast by proteolytic cleavage that regulate immune and stress responses ^22^. PEPSI6 localizes to root plastids and engages plastid import and proteostasis machinery in a PrLD-dependent manner. Quantitative proteomics revealed that PEPSI6 specifically increases the abundance of the plastidial enzyme 1-deoxy-D-xylulose 5-phosphate reductoisomerase (*At*DXR), which catalyzes the first committed step of the methylerythritol phosphate (MEP) pathway for isoprenoid biosynthesis ^23^. During root colonization, PEPSI6 also undergoes partial apoplastic proteolytic cleavage of the CAP domain, revealing a PrLD-independent activity. Both full-length and PrLD-deficient PEPSI6 enhance fungal colonization, indicating separable plastidial and apoplastic contributions to symbiosis. Together, these findings identify DXR as an effector target and reveal PrLDs as noncanonical plastid-targeting elements through which mutualistic fungi modulate plastid metabolism.

## Results

### A class of PrLD-containing effector proteins (PEPSIs) is induced during *S. indica* root colonization

Prion-like domains (PrLDs) can modulate protein behavior by influencing localization, interaction networks, trafficking, phase transitions, and condensate formation ^24^. Our recent work showed that exogenous non-plant proteins containing PrLDs can be imported into chloroplasts ^19^, suggesting that PrLDs, or prion-like regions within intrinsically disordered regions (IDRs), may engage plastid import machinery. Based on these findings, we searched the *S. indica* secretome for PrLD-containing effectors that might play specialized roles during root colonization. We first predicted secreted proteins using Predector ^25^ and then screened the *S. indica* proteome for PrLDs using the PLAAC algorithm ^26^. To refine the candidate list, we integrated available transcriptomic datasets and selected genes consistently upregulated during *Arabidopsis* root colonization at different time points (**Fig. 1**). This pipeline identified 16 PrLD-containing secreted proteins, which we termed PEPSIs. These candidates spanned diverse predicted functional domains, including several glycoside hydrolases ^27^, four predicted to have esterase activity, and one containing a CAP domain (**Fig. 1**).

**Figure 1.**
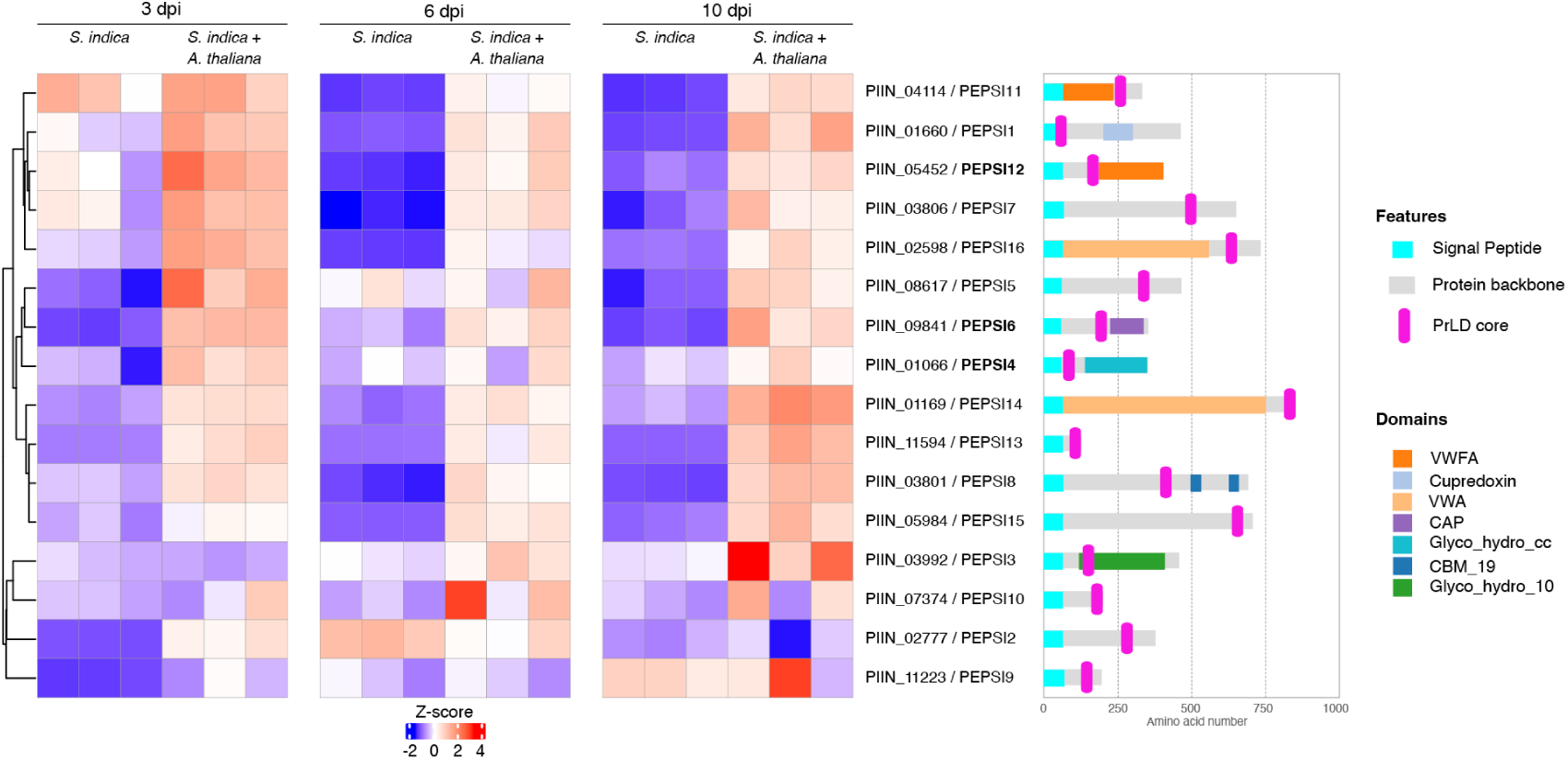
Identification of prion-like domain–containing effector genes from *S. indica* induced during plant interaction. **Left**, heatmap showing row-wise z-scored transcript abundance (TPM) of prion-like effector protein (PEPSI) candidates in *S. indica* grown alone or in the presence of *Arabidopsis thaliana* roots at 3, 6, and 10 days post inoculation (dpi). Genes were hierarchically clustered using the ComplexHeatmap package. Data represent three biological replicates per condition. PIIN identifiers are shown, and the three effectors selected for further characterization (PEPSI4, PEPSI6, and PEPSI12) are indicated in bold. **Right**, schematic representation of PEPSI domain architectures. Signal peptides are shown in cyan, protein backbone in gray, PrLD cores in magenta, and additional predicted domains are shown as colored rectangles.

Given their induction during colonization, we hypothesized that these PEPSIs contribute to symbiosis establishment. For further characterization, we selected PEPSI4 (PIIN_01066), PEPSI6 (PIIN_09841), and PEPSI12 (PIIN_05452), representing distinct domain architectures (**Fig. 1**).

### PrLDs are required for plastid targeting of three PEPSI effectors

To examine the subcellular localization of selected PEPSIs, we synthesized codon-optimized versions of PEPSI4, PEPSI6, and PEPSI12, including their signal peptides, verified their amino-acid sequences against transcriptomic data (**Extended Data Fig. 1**) and fused them to GFP for transient expression in *Nicotiana benthamiana* (**Fig. 2a**). Upon agroinfiltration, control GFP showed the expected cytoplasmic and nuclear distribution, whereas GFP-tagged PEPSI4, PEPSI6, and PEPSI12 exhibited distinct localization patterns (**Fig. 2b**). Notably, PEPSI4 and PEPSI6 formed discrete cytoplasmic speckles, consistent with the propensity of PrLD-containing proteins to form biomolecular condensates. Strikingly, all three PEPSIs also accumulated within chloroplasts, with PEPSI4 additionally forming speckles adjacent to chloroplasts, possibly reflecting plastid-proximal assemblies (**Fig. 2b**) ^28^.

**Figure 2.**
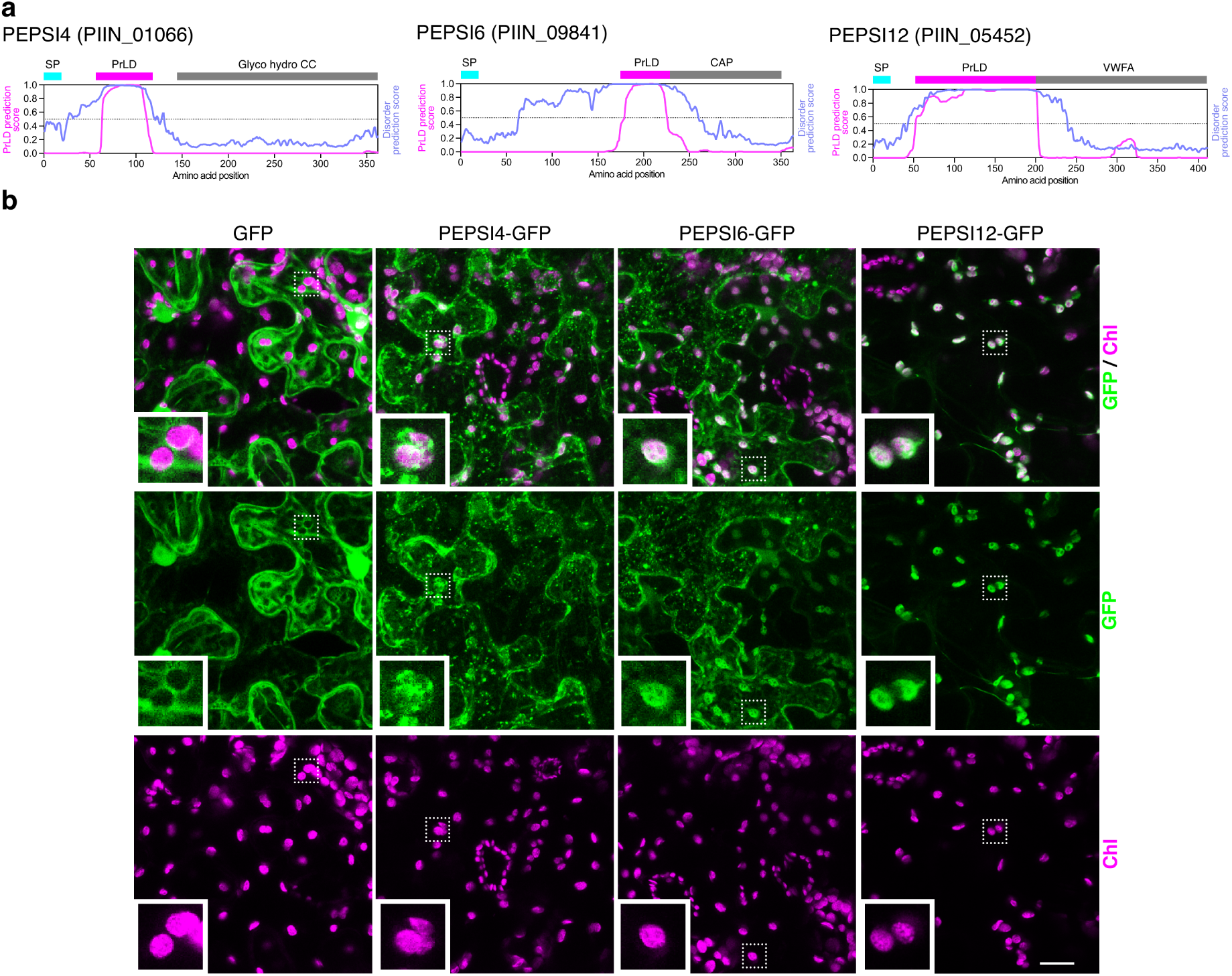
Subcellular localization of selected *S. indica* prion-like effector proteins in *N. benthamiana*. **a**, Schematic representation of PEPSI4, PEPSI6, and PEPSI12 showing prediction profiles for prion-like domains (PrLDs) and intrinsically disordered regions (IDRs). Cyan boxes indicate predicted signal peptides (SP), magenta boxes denote PrLDs, and gray boxes represent additional predicted Pfam domains. Blue traces indicate IDR prediction scores, and magenta traces indicate PrLD prediction scores along the protein sequence. **b**, Representative confocal microscopy images of *N. benthamiana* leaf epidermal pavement cells transiently expressing GFP alone or GFP-tagged PEPSI4, PEPSI6, or PEPSI12, each including its native N-terminal signal peptide (SP), under control of the 35S promoter, imaged 3 days post agroinfiltration. GFP fluorescence is shown in green and chloroplast autofluorescence in magenta. Scale bar, 20 µm. Images are representative of three independent experiments.

High-magnification imaging further confirmed chloroplast accumulation of PEPSI4, PEPSI6, and PEPSI12 (**Fig. 3a**). As a positive control for plastid targeting, we expressed GFP fused to the transit peptide of the plastid-localized heat shock protein *At*HSP21/AT4G27670 (pGFP) (**Fig. 3a; Extended Data Fig. 2**). The consistent plastid localization of all three effectors suggested that PrLDs might contribute to noncanonical plastid targeting. To test this, we generated truncated versions lacking their prion-like domains (PEPSI4ΔP, PEPSI6ΔP, and PEPSI12ΔP) (**Fig. 3b**). Upon agroinfiltration, all PrLD-deleted constructs lost chloroplast localization, and the condensate-like speckles characteristic of the full-length proteins were no longer observed (**Fig. 3c; Extended Data Fig. 3**), demonstrating that PrLDs are required for both speckle formation and plastid targeting.

**Figure 3.**
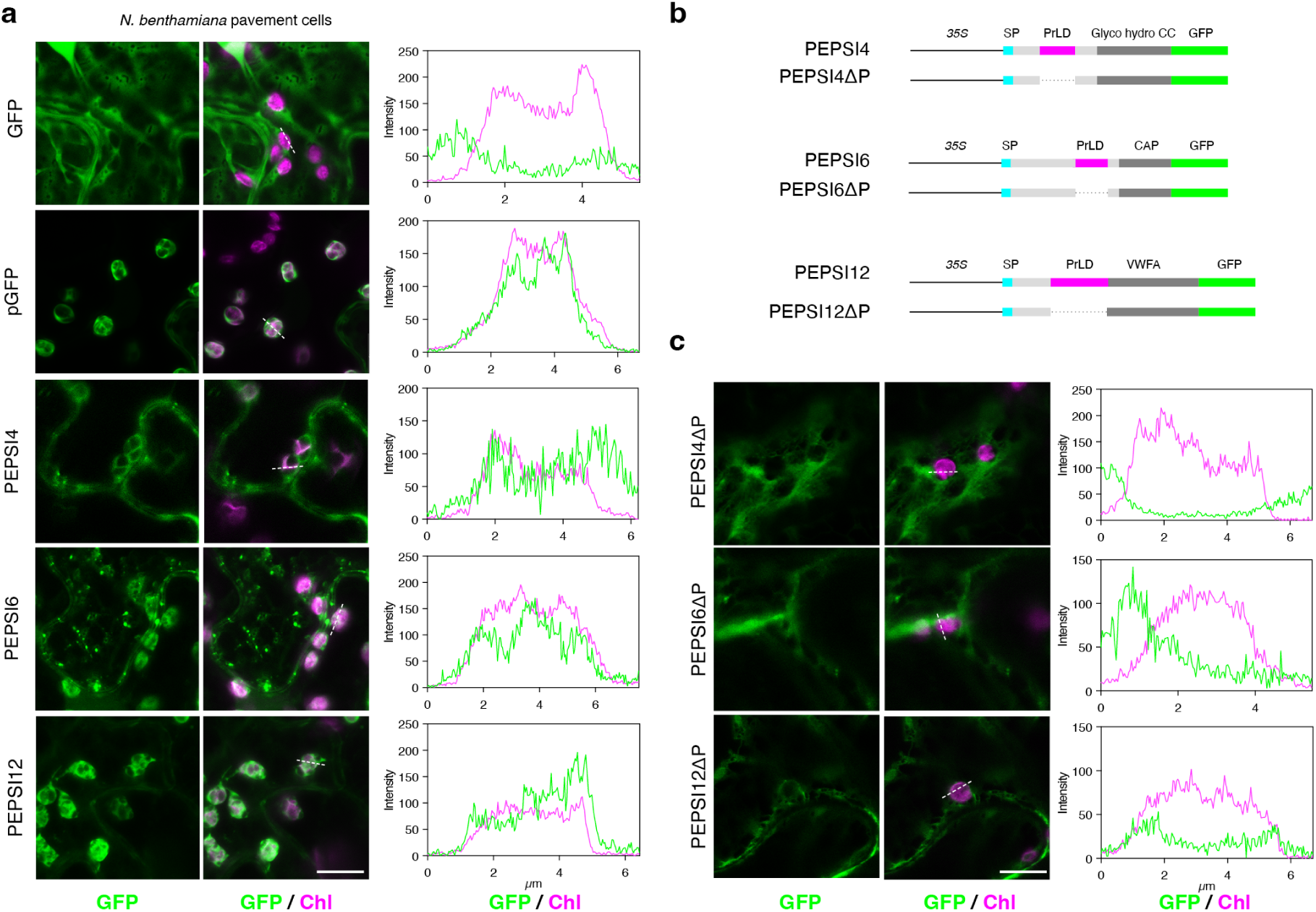
Prion-like domains are required for plastid localization of PEPSI4, PEPSI6 and PEPSI12 effectors. **a**, Representative confocal microscopy images of *Nicotiana benthamiana* leaf epidermal pavement cells transiently expressing C-terminal GFP-tagged PEPSI4, PEPSI6, or PEPSI12, each including its native N-terminal signal peptide (SP), under control of the 35S promoter. A plastid-targeted GFP construct (pGFP) was included as a positive control, and cytosolic GFP served as a negative control. A secretion control is shown in Fig. 6d. GFP fluorescence is shown in green and chloroplast autofluorescence in magenta. Line plots depict GFP and chlorophyll fluorescence intensity profiles measured along the indicated dashed lines. Scale bar, 10 µm. **b**, Schematic representation of PEPSI domain architecture, showing predicted signal peptides (SP), prion-like domains (PrLD, magenta), additional predicted functional domains (gray), and the regions deleted in the corresponding PrLD-deletion (PEPSIΔP) variants. **c**, Representative confocal microscopy images of *N. benthamiana* epidermal cells expressing C-terminal GFP-tagged PEPSIΔP variants. GFP fluorescence (green) and chloroplast autofluorescence (magenta) are shown. Line plots depict fluorescence intensity profiles measured along the indicated dashed lines. Scale bar, 10 µm. All images are representative of three independent experiments.

### PEPSI6 and PEPSI12 accumulate in plastids in Arabidopsis root cells and enhance fungal colonization

To test whether plastid-associated accumulation of PEPSI candidates also occurs in root tissue, we generated stable Arabidopsis transgenic lines expressing PEPSI6–GFP, including its native fungal signal peptide (**Extended Data Fig. 4**), along with a plastid-targeted GFP control line (pGFP) (**Extended Data Fig. 2**). Confocal imaging of multiple root regions revealed that PEPSI6 accumulates in plastid-like structures, showing localization patterns similar to pGFP (**Fig. 4a,b**). In contrast, GFP control plants displayed the expected cytosolic and nuclear fluorescence (**Fig. 4a**).

**Figure 4.**
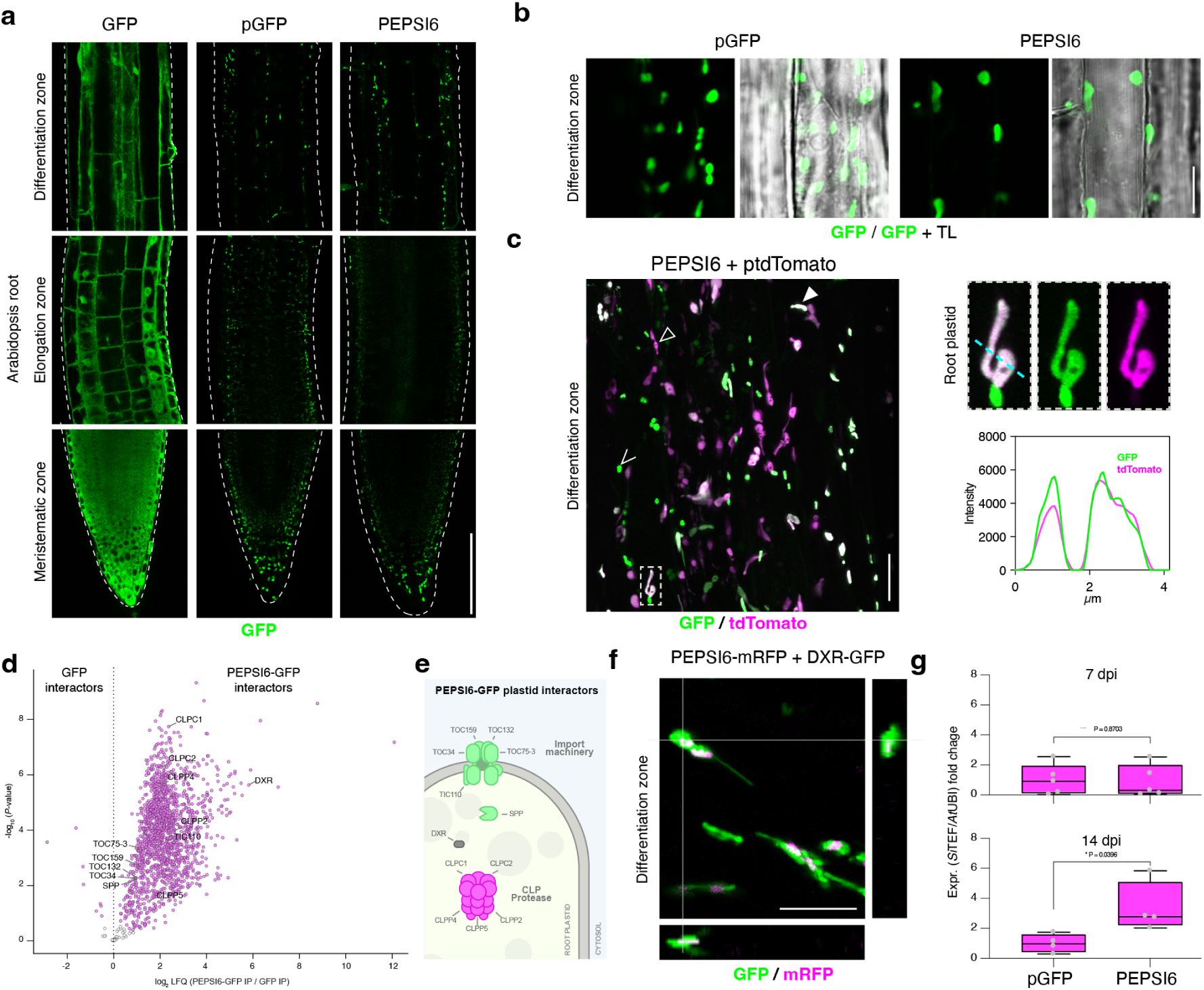
PEPSI6 localizes to plastids in Arabidopsis root cells and associates with DXR. **a**, Representative confocal microscopy images of 7-day-old *Arabidopsis thaliana* roots expressing PEPSI6 fused to C-terminal GFP. A cytosolic GFP control and a plastid-targeted GFP construct (pGFP) were included for comparison. Images show single optical sections of the meristematic, elongation, and differentiation zones of the root. Scale bar, 100 µm. **b**, Magnified views of cells in the differentiation zone of roots expressing pGFP or PEPSI6–GFP. GFP fluorescence is shown alone and overlaid with transmitted light (TL). Scale bar, 10 µm. Panels **a** and **b** are representative of three independent roots. **c**, Representative confocal microscopy image of an Arabidopsis root co-expressing PEPSI6–GFP and a plastid-targeted tdTomato marker (ptdTomato). Z-projection of 183 z-stacks spanning a total depth of 36.4 µm and merged GFP and tdTomato fluorescence. Open arrowheads indicate PEPSI6-containing puncta outside plastids, closed white arrowheads indicate colocalization of PEPSI6 with plastids, and closed black arrowheads mark plastids lacking detectable PEPSI6–GFP signal. Insets show individual plastids and corresponding GFP and tdTomato fluorescence intensity profiles along the indicated line. Scale bar, 20 µm. **d**, Volcano plot of proteins co-immunoprecipitated with PEPSI6–GFP from 7-day-old Arabidopsis roots, plotted as log₂ fold change in label-free quantification (LFQ) intensity versus –log₁₀(P value) relative to a cytosolic GFP control. Magenta dots indicate significantly enriched proteins after multiple-testing correction (two-tailed Student’s t test; n = 4 biological replicates; FDR-adjusted q value < 0.05). **e**, Schematic representation of selected PEPSI6–GFP interactors localized to root plastids, including components of the plastid import machinery, proteostasis pathways, and DXR. **f**, Representative confocal microscopy image of an Arabidopsis root co-expressing PEPSI6–mRFP and plastid-localized DXR–GFP. Orthogonal views of 25 z-stacks spanning a total depth of 5 µm and merged fluorescence channels confirm colocalization within plastids. Scale bar, 10 µm. **g**, Intraradical *S. indica* colonization quantified by qPCR in seed-inoculated pGFP and PEPSI6–GFP plants at 7 and 14 days post inoculation (dpi). Values are normalized to pGFP. Statistical analysis was performed using a two-tailed Student’s t test for unpaired samples.

To validate plastid localization, we introduced a plastid-targeted tdTomato marker (ptdTomato) into the PEPSI6–GFP line (**Extended Data Fig. 5**). Confocal microscopy showed extensive co-localization of PEPSI6–GFP with ptdTomato, confirming plastid targeting in roots (**Fig. 4c; Supplementary Video 1**). A subset of GFP-positive puncta did not co-localize with ptdTomato, suggesting the presence of additional PEPSI6-containing assemblies outside plastids (**Fig. 4a**), similar to cytoplasmic speckles observed in *N. benthamiana* epidermal cells. These puncta exhibited highly dynamic movement (**Supplementary Video 1**), consistent with transient or mobile protein assemblies.

To identify host proteins interacting with PEPSI6 *in planta*, we performed co-immunoprecipitation followed by label-free proteomics using roots expressing PEPSI6–GFP and cytosolic GFP as a control. The PEPSI6–GFP pulldown was significantly enriched for plastid-localized proteins associated with protein import, metabolism, and proteolysis (**Fig. 4d,e; Extended Data Fig. 6; Supplementary Table 1**). These included components of the plastid import machinery TOC/TIC translocon, subunits of the CLP protease complex, and the stromal processing peptidase SPP. Notably, PEPSI6 also co-purified with DXR, a plastid-resident enzyme of the MEP pathway.

To further examine this association, we performed confocal co-localization analyses *in planta* by expressing PEPSI6–mRFP in transgenic plants carrying plastid-localized DXR–GFP ^29^. Imaging revealed co-localization of PEPSI6 with DXR in plastids (**Fig. 4f**), supporting their association in this compartment.

To assess the biological relevance of PEPSI6 during colonization by *S. indica*, we quantified intraradical fungal biomass by qPCR in roots of pGFP and PEPSI6–GFP plants at 7 and 14 days post inoculation (dpi). While no significant differences were observed at 7 dpi, PEPSI6–GFP lines showed significantly increased colonization at 14 dpi compared to controls (**Fig. 4g**), indicating that PEPSI6 promotes fungal colonization at later stages.

Finally, to determine whether this plastid-associated accumulation extends to other PEPSI members, we generated stable Arabidopsis lines expressing PEPSI12–GFP. Similar to PEPSI6, PEPSI12 accumulated in plastid-like structures in roots, and PEPSI12-expressing lines also showed enhanced *S. indica* colonization at 14 dpi (**Extended Data Fig. 7**). Together, these results show that two PEPSI proteins with distinct domain architectures but shared IDR-embedded PrLDs localize to plastids in root tissue and promote fungal colonization.

### PrLD-dependent plastid targeting enables PEPSI6 to associate with DXR and alter MEP-pathway metabolism

We next tested whether PrLD deletion also impaired plastid targeting of PEPSI6 in *Arabidopsis* roots. Unlike full-length PEPSI6–GFP, PEPSI6ΔP–GFP did not localize to root plastids in stable transgenic plants (**Fig. 5a–c; Extended Data Fig. 8**). These results show that the PrLD is required for plastid localization.

**Figure 5.**
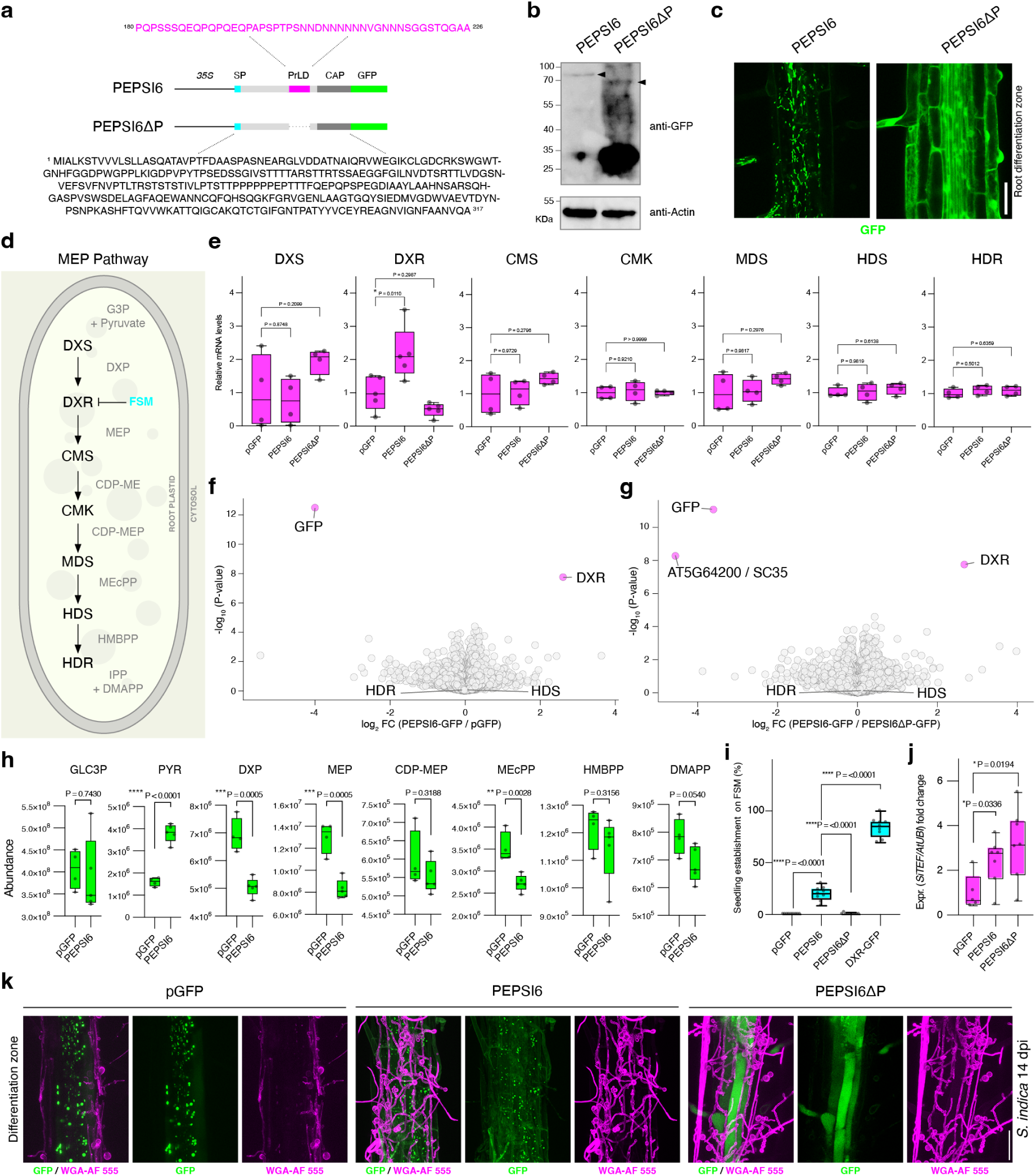
The prion-like domains is required for PEPSI6 plastid targeting and DXR modulation in roots. **a**, Schematic representation of PEPSI6 constructs showing predicted domains. Cyan boxes indicate signal peptides (SP), magenta boxes denote prion-like domains (PrLD), dark gray boxes represent predicted functional domains, and green boxes represent GFP. The PrLD amino acid sequence is shown in magenta, and the corresponding sequence of the construct lacking the PrLD is also displayed. **b**, Immunoblot analysis of 14-day-old root tissue expressing PEPSI6–GFP or PEPSI6ΔP–GFP, detected with an anti–GFP antibody. Actin was used as a loading control. Black arrowheads indicate the expected molecular sizes of PEPSI6–GFP (∼76 kDa) and PEPSI6ΔP–GFP (∼64 kDa). Representative of two independent experiments. **c**, Representative confocal microscopy images of Arabidopsis roots expressing PEPSI6–GFP or PEPSI6ΔP–GFP. Images show the GFP channel. Scale bar, 50 µm. **d**, Schematic representation of the plastidial methylerythritol phosphate (MEP) pathway, showing associated enzymes, the inhibitor fosmidomycin (FSM), and metabolites. **e**, qPCR analysis of MEP pathway gene expression in pGFP control plants and PEPSI6 transgenic lines. **f, g**, Volcano plots comparing the root proteomes of (**f**) PEPSI6–GFP versus pGFP and (**g**) PEPSI6–GFP versus PEPSI6ΔP–GFP in 20-day-old Arabidopsis roots. Plots show –log₁₀(P value) versus log₂ fold change of protein label-free quantification (LFQ) intensities. Magenta dots indicate significantly changed proteins (two-tailed Student’s t test; n = 4–5 biological replicates). **h**, Box plots showing the abundance of metabolites from the MEP pathway and associated precursors in pGFP and PEPSI6–GFP roots under mock conditions, measured by LC–MS/MS. Statistical significance was assessed using a two-tailed unpaired Student’s t test. **i**, Seedling establishment on medium supplemented with FSM, expressed as the percentage of seedlings developing green cotyledons and true leaves. pGFP, PEPSI6–GFP, PEPSI6ΔP–GFP, and DXR–GFP lines are shown. These results were corroborated by pulse-amplitude modulated (PAM) chlorophyll fluorescence measurements, yielding similar results. **j**, Intraradical *S. indica* colonization quantified by qPCR in seed-inoculated pGFP, PEPSI6–GFP, and PEPSI6ΔP–GFP plants at 14 days post inoculation (dpi). Values are expressed relative to pGFP. Statistical analysis was performed using a two-tailed Student’s t test for unpaired samples. **k**, Representative confocal microscopy images of colonized roots from the lines described in **j**. *S. indica* hyphae were visualized using WGA–Alexa Fluor 555 staining.

Because PEPSI6 accumulates within plastids and interacts with DXR, we examined whether it alters the MEP pathway, which is modulated during *S. indica* colonization ^16^ (**Fig. 5d**). We analyzed MEP gene expression and plastid metabolites in PEPSI6–GFP and control plants. Among all enzymes tested, only DXR showed increased transcript abundance in PEPSI6–GFP plants, whereas PEPSI6ΔP–GFP plants resembled the pGFP control (**Fig. 5e**).

To assess PEPSI6-dependent changes at the protein level, we performed label-free quantitative proteomics on roots expressing PEPSI6–GFP, PEPSI6ΔP–GFP, and pGFP. Direct pairwise comparisons of PEPSI6–GFP versus pGFP and PEPSI6–GFP versus PEPSI6ΔP–GFP identified DXR as the sole protein specifically enriched in the presence of full-length PEPSI6 (**Fig. 5f,g; Supplementary Table 2**). These data indicate that DXR accumulation is specifically associated with full-length PEPSI6 expression and requires its PrLD.

Consistent with this protein-level specificity, metabolite profiling by liquid chromatography–tandem mass spectrometry (LC-MS/MS) revealed decreases in early MEP-pathway intermediates, including DXP, MEP, and MEcPP, while downstream products such as HMBPP remained unchanged (**Fig. 5h**).

Because DXR accumulation and PEPSI6 plastidial localization depended on the presence of the PrLD, we next examined how its removal alters host protein associations. To this end, we compared the interactomes of PEPSI6–GFP and PEPSI6ΔP–GFP using co-immunoprecipitation and label-free proteomics, as described above. PEPSI6ΔP–GFP lacked plastid-associated interactors (**Extended Data Fig. 9; Supplementary Table 1**), indicating that the PrLD is required for PEPSI6 to access plastid-associated protein networks and associate with DXR.

To assess whether PEPSI6 modulates plant responses to perturbation of the DXR step in the plastidial MEP pathway, *Arabidopsis* seeds were germinated on medium supplemented with fosmidomycin (FSM), a specific inhibitor of DXR in plants and bacteria, or on mock medium. FSM treatment causes seedling growth arrest and cotyledon bleaching; under these conditions, seedling establishment can be quantified as the proportion of seedlings that successfully green and develop true leaves ^30^. Notably, PEPSI6 overexpressing plants exhibited increased tolerance to FSM, as quantified by a significantly higher seedling establishment rate, whereas the PEPSI6ΔP line remained highly sensitive, comparable to the pGFP control. DXR–GFP overexpressing plants served as a tolerant reference (**Fig. 5i**).

We next examined whether PrLD-dependent plastid targeting of PEPSI6 is required for enhanced fungal colonization. To this end, we quantified intraradical *S. indica* colonization by qPCR at 14 dpi in pGFP, PEPSI6–GFP, and PEPSI6ΔP–GFP lines. Both PEPSI6 variants exhibited increased root colonization compared with the pGFP control, indicating that PEPSI6 promotes fungal colonization independently of its PrLD-mediated plastid targeting (**Fig. 5j**). Visualization of fungal structures using WGA–AF555 confirmed these trends for extraradical colonization (**Fig. 5k**). Thus, while PrLD-dependent plastid targeting enables PEPSI6 to associate with DXR and alter MEP-pathway metabolism, enhanced colonization does not require the PrLD, indicating a plastid-independent function of this effector.

### PEPSI6 undergoes partial apoplastic processing during colonization

Western blot analysis of PEPSI6–GFP plants revealed a lower–molecular weight GFP-tagged fragment that increased upon *S. indica* colonization and was only faintly detectable in mock-treated roots (**Fig. 6a**). This colonization-associated fragment is consistent with proteolytic cleavage of PEPSI6 *in planta*, likely within its C-terminal region. Structural analysis revealed that, in addition to the intrinsically disordered region and PrLD, PEPSI6 contains a C-terminal CAP domain structurally similar to that of the *Arabidopsis* pathogenesis-related protein PR1 (**Fig. 6b; Extended Data Fig. 10**).

**Figure 6.**
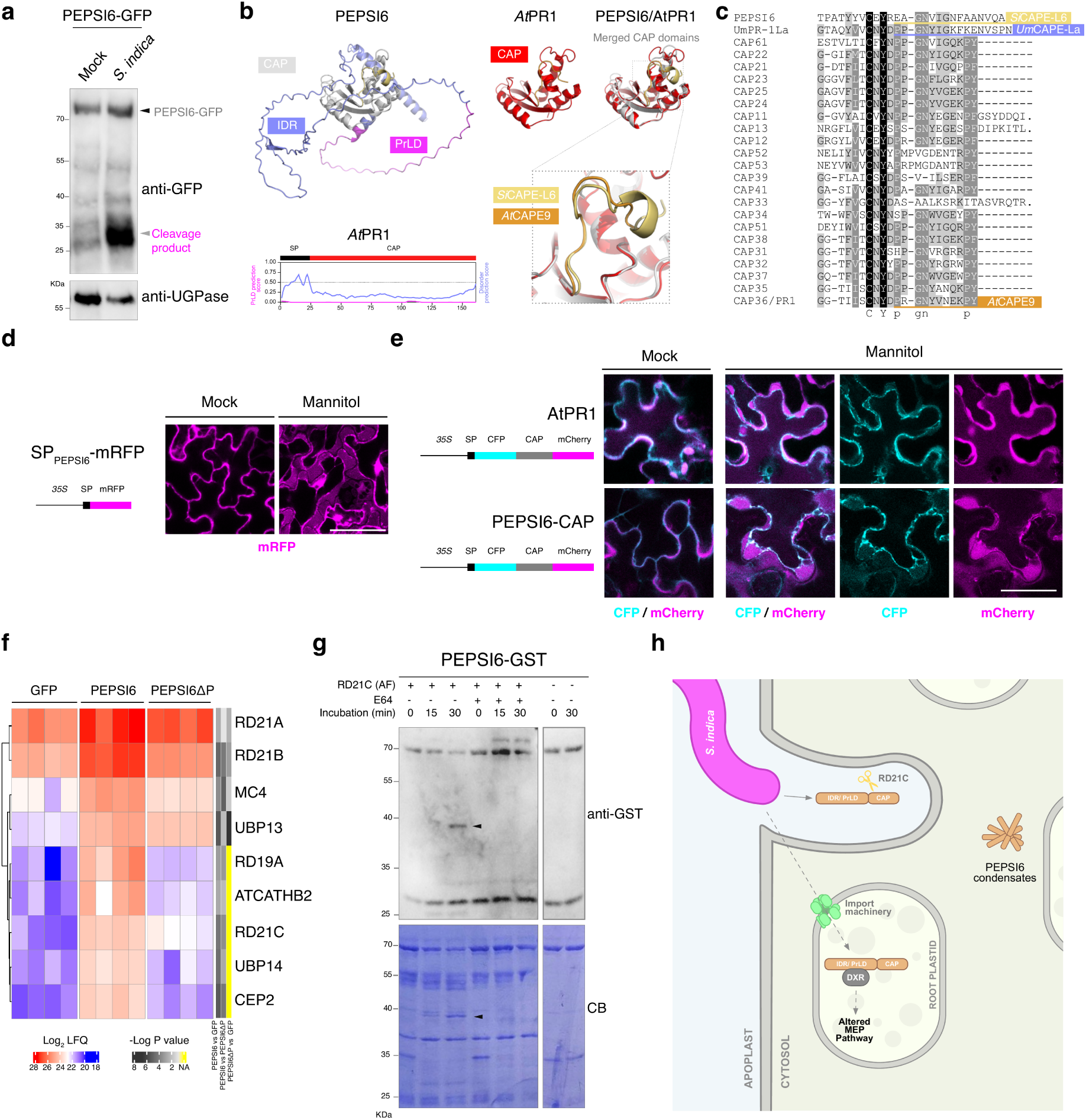
PEPSI6 undergoes C-terminal proteolytic processing and harbors a PR1-like CAP domain. **a**, Immunoblot analysis of root tissue from 14-day-old plants expressing PEPSI6–GFP under mock or *S. indica*–colonized conditions, detected using an anti–GFP antibody. UGPase was used as a loading control. The black arrowhead indicates the expected molecular weight of full-length PEPSI6–GFP, whereas the gray arrow indicates a cleavage product that accumulates upon *S. indica* colonization. The blot is representative of three independent experiments. **b**, AlphaFold structural predictions of *S. indica* PEPSI6 and *A. thaliana* PR1, with signal peptides excluded from the models. A schematic representation of PR1 domain architecture is shown, in which no intrinsically disordered region (IDR) or prion-like domain (PrLD) is detected. The N-terminal signal peptide (SP) is indicated by a black box and the CAP domain by a red box. Disorder scores (IUPred2A) are shown in lilac and PrLD prediction scores (PLAAC) in magenta. **c**, Multiple sequence alignment of the C-terminal regions of selected CAP proteins from *A. thaliana* ^57^, *S. indica*, and *Ustilago maydis* (UMAG_01204) ^31^, with the putative C-terminal CAP-derived peptide (CAPE) region indicated. **d**, Representative confocal microscopy images show apoplastic localization of mRFP fused to the native PEPSI6 signal peptide following transient expression in *N. benthamiana* leaf epidermal pavement cells. Leaves were treated with mock solution or 0.8 M mannitol for 1 h prior to imaging. mRFP fluorescence is shown in magenta. Scale bar, 50 µm. **e**, Subcellular localization of double-tagged Arabidopsis PR1 and the PEPSI6 CAP region. Schematics depict the constructs indicating the signal peptide (black), CFP (cyan), CAP domain (gray), and mCherry (magenta). Samples were treated with mock solution or 0.8 M mannitol prior to visualization. Scale bar, 50 µm. **f**, Heat map showing log₂-transformed LFQ intensities of proteins co-immunoprecipitated with GFP from pGFP, PEPSI6–GFP, and PEPSI6ΔP–GFP expressing roots, highlighting proteins annotated with cysteine protease activity. The –log₁₀(P value) bar indicates statistical significance between comparisons. **g**, *In vitro* cleavage assays of PEPSI6–GST incubated over time with heterologously expressed RD21C. Cleavage products were detected by immunoblotting using an anti-GST antibody, and total protein loading is shown by Coomassie Brilliant Blue staining (CB). The papain-like cysteine protease inhibitor E64 was used to block PLCP activity. **h**, Proposed model illustrating two PEPSI6 populations: one localized to the apoplast and subject to proteolytic processing by papain-like cysteine proteases such as RD21C, and a second population targeted to plastids, where PEPSI6 interacts with DXR and modulates MEP-pathway metabolism.

CAP-domain proteins from plants and fungi can undergo apoplastic processing to release short bioactive peptides with immune-modulatory functions. In *Arabidopsis*, PR1 is cleaved by the papain-like cysteine protease XCP1 at the caspase-like CNYD motif during salicylic acid-induced immunity, releasing the CAP-derived peptide CAPE9 ^22^. Similarly, the maize pathogen *Ustilago maydis* secretes the PR1-like effector *Um*PR-1La, which is processed by the host papain-like cysteine protease CatB3 at the conserved CNYD site to yield a CAPE-like peptide that subverts plant defenses and promotes virulence ^31^. Although PEPSI6 lacks the canonical CNYD motif shared by PR1 and *Um*PR-1La, it carries a related C-terminal CXYX sequence (**Fig. 6c; Extended Data Fig. 10**), raising the possibility of cleavage by a distinct protease repertoire encountered during microbial interactions.

To test whether the CAP region of PEPSI6 reaches the extracellular space, we generated a double-tagged secreted reporter in which the PEPSI6 signal peptide and CFP were fused to the N terminus and mCherry to the C terminus of the CAP region. As a positive control for secretion, we expressed an SP_PEPSI6_-mRFP fusion, in which the native PEPSI6 signal peptide was directly fused to mRFP. This construct accumulated in the apoplast of *N. benthamiana* epidermal cells and became clearly visible after plasmolysis, supporting recognition of the fungal signal peptide and efficient secretion in plant cells (**Fig. 6d**). We then examined the subcellular distribution of the double-tagged CAP reporter. Following plasmolysis, the C-terminal mCherry signal remained in the apoplast (**Fig. 6e**), resembling the behavior of Arabidopsis PR1 analyzed in parallel (**Fig. 6e**).

To identify plant proteases potentially associated with PEPSI6 cleavage, we analyzed the chitohexaose-activated PEPSI6–GFP interactome from roots. Although protease–substrate interactions are often transient and difficult to capture by co-immunoprecipitation, several proteases were significantly enriched relative to cytosolic GFP controls, with a subset also detected in the PEPSI6ΔP interactome **(Fig. 6f; Supplementary Table 1)**. Notably, the canonical PR1-processing protease XCP1 was not detected, whereas other papain-like cysteine proteases expressed in roots, including RD21A and RD21C, were present (**Fig. 6f**). This pattern is consistent with the protease landscape of root tissues and supports the idea that PEPSI6 processing involves a protease repertoire distinct from the canonical PR1-associated pathway.

To test whether apoplastic fluid containing active RD21 proteases can cleave PEPSI6 directly, we incubated recombinant His–PEPSI6–GST with apoplastic fluid isolated from *N. benthamiana* leaves overexpressing RD21C (**Fig. 6g**). We confirmed that RD21C is active in the apoplast by activity-based protein profiling, whereas the RD21A catalytic mutant used as a control lacked detectable activity (**Extended Data Fig. 11**). Incubation with RD21C-containing apoplastic fluid reproducibly yielded a discrete lower–molecular weight fragment, whereas no cleavage of His–PEPSI6–GST was detected in the presence of apoplastic fluid from the catalytic mutant. Cleavage was suppressed by the papain-like cysteine protease inhibitor E64 (**Fig. 6g**), indicating that cleavage depends on active papain-like cysteine protease activity in RD21C-containing apoplastic fluid. The size of the detected fragment was larger than expected for a CAPE-like peptide fused to GST, suggesting that RD21C-containing apoplastic extracts mediate an initial cleavage of PEPSI6, while additional proteolytic steps may be required to generate smaller peptides.

Together, these findings are consistent with a model in which PEPSI6 is secreted into the apoplast, where a fraction undergoes proteolytic processing by host proteases, whereas uncleaved PEPSI6 may enter host cells and be imported into plastids. This apparent dual fate places PEPSI6 in spatially distinct host compartments and is consistent with its PrLD-dependent association with DXR and its contribution to mutualistic root colonization (**Fig. 6h**).

## Discussion

Our study reveals a previously unrecognized role for PrLDs embedded in intrinsically disordered regions in promoting the accumulation of fungal effectors in plant plastids. We show that a subset of *S. indica* secreted effectors, termed PEPSIs, localize in root plastids in a PrLD-dependent manner, and that one member, PEPSI6, measurably alters plastid-associated molecular outputs. Together, these findings identify PrLDs as potential noncanonical elements that can facilitate plastid accumulation in plants. The enrichment of TOC/TIC components, stromal processing factors such as SPP, subunits of the CLP protease system, and other plastid proteostasis modules in the PEPSI6 interactome is consistent with PEPSI6 engaging host pathways that normally regulate import and quality control of nuclear-encoded plastid proteins. A parsimonious model is that PrLD-dependent interactions increase the likelihood that PEPSI6 is routed into, or retained within, the plastid proteostasis environment, thereby enhancing plastid accumulation. Quantitative proteomics and gene expression analyses consistently identified DXR as the only plastidial protein upregulated in PEPSI6-expressing plants. Physical association, plastid co-localization, and metabolite profiling further support DXR as an effector target. Metabolite profiling showed that the early intermediates DXP and MEP decreased relative to controls, whereas downstream products remained unchanged. This pattern indicates that PEPSI6 shifts steady-state levels of selected metabolites within the MEP pathway. Because DXP and MEP lie immediately upstream and downstream of DXR, respectively, they are expected to respond most directly to changes in DXR abundance, activity, or stability, while distal metabolites such as HMBPP can remain buffered by homeostatic regulation. This aligns with a fine-tuned modulation of plastid metabolism and retrograde signaling, for example through reduced MEcPP levels, rather than broad alteration of isoprenoid biosynthesis. During *S. indica* colonization, additional plastid-targeted effectors and/or stress-dependent demands on the pathway may also shape MEP-pathway outputs and contribute to stronger metabolic changes than those observed under our experimental conditions.

Notably, PEPSI6-mediated modulation of DXR parallels the induction of DXR reported during arbuscular mycorrhizal symbiosis, where DXR transcripts and protein accumulate specifically in arbuscule-containing cells. In maize roots, DXR localizes to a dense network of plastids and stromules surrounding developing arbuscules and reaches maximal abundance during arbuscule maturity and early senescence ^6^. This spatially restricted increase has been proposed to support localized MEP flux and apocarotenoid-related metabolism associated with arbuscule accommodation. More broadly, recent work shows that an unrelated *S. indica* effector, *Si*E141, perturbs plastid protein targeting by binding the thioredoxin-like plastid protein CDSP32 and redirecting it to the nucleus, thereby reshaping host stress and immune outputs ^5^. Together with our findings, these observations support the view that plastids contribute to the cellular reprogramming required for beneficial root–fungus interactions and may therefore represent recurrent targets of fungal effectors. An important next step will be to define the collective function of this putative plastid-targeting effector network during colonization.

Beyond plastid targeting, the PrLD-containing effector PEPSI6 assembles into dynamic cytoplasmic puncta in both *N. benthamiana* leaf and Arabidopsis root cells. Given the intrinsic disorder and Q/N-rich composition of its PrLD, these puncta are consistent with higher-order assemblies and may represent biomolecular condensates. Phase separation is increasingly recognized as an organizing principle in plant stress signaling and immunity ^32^, with intrinsically disordered regions enabling rapid, reversible interactions with highly connected host factors ^33^. In this context, pathogen effectors enriched in IDRs or low-complexity regions may exploit similar multivalent interaction modes to access central host regulatory nodes ^21^. Formation of these puncta requires the PrLD. One possibility is that PEPSI6 forms transient assemblies resembling import-competent precursor proteins or excess plastid-destined substrates, which could be sensed by plastid-associated quality-control systems that monitor protein import and proteostasis ^34^. Chloroplast protein homeostasis is tightly coupled to immunity and involves chaperones, proteases, and retrograde signaling components that surveil proteome integrity and stress status ^35^. Engagement of these pathways by PEPSI6 could facilitate plastid entry, stabilize the effector post-import, or modulate defense-relevant signaling outputs.

In addition to its plastid-targeted activity, PEPSI6 carries a CAP domain that undergoes partial proteolytic processing during root colonization. Our results show that the CAP region can accumulate in the apoplast, consistent with proteolytic processing *in planta*. Protease interactome analyses and *in vitro* assays point to papain-like cysteine proteases, including members of the RD21 subfamily, as candidates capable of mediating PEPSI6 cleavage and generating a CAP-derived fragment in root tissues. Whether this fragment is further processed into a CAPE-like peptide remains to be determined. This processing mechanism differs from the canonical PR1 pathway mediated by XCP1 and instead may reflect the distinct protease environment of root tissues. PEPSI6 interaction with and processing by RD21-family proteases may intersect with the broader activity of host papain-like cysteine proteases. RD21s have emerged as central protease hubs in plant–microbe interactions and are recurrently targeted by diverse pathogen-secreted effectors, not only for enzymatic inhibition but also for sequestration, functional hijacking, mislocalization, or proteolytic destabilization ^36^.

Notably, both full-length PEPSI6 and the plastid-targeting–deficient PEPSI6ΔP enhance *S. indica* colonization, indicating that PEPSI6 contributes to symbiosis through more than one functional route. One activity depends on the PrLD and is associated with plastid targeting, DXR engagement, and modulation of plastid metabolism, whereas a second contribution to colonization does not require the PrLD and is therefore likely plastid independent. This dual function is consistent with the multifunctionality seen in other effectors and underscores how a single fungal protein can coordinate processes across multiple host compartments to fine-tune host physiology during symbiosis.

Together, these findings highlight plastids as important targets of mutualistic fungal effectors, expand the repertoire of noncanonical domain architectures associated with plastid access, and illustrate how fungal effectors can integrate spatially separated and functionally distinct activities.

## Material and methods

### Identification of prion-like domain-containing *S. indica* effectors induced during plant colonization

To identify prion-like domain-containing (PrLD) effectors, the *Serendipita indica* proteome was scanned using PLAAC ^26^ with *S. indica*–specific background amino acid probabilities (-a 0) and a sliding window size of 40 amino acids (-c 40). As previously described ^33^, proteins with a COREscore > 0 were considered to have a high probability of containing a PrLD. In parallel, the proteome was analyzed using Predector ^25^ to identify putative secreted effector proteins, and only proteins predicted to be secreted (is_secreted = 1) were retained. These candidates were further filtered based on our transcriptomic data, selecting proteins that were consistently upregulated during *S. indica in planta* interactions at a minimum of two time points. Gene expression data were obtained from the resequencing of the *Serendipita indica* transcriptome dataset previously available ^15,37^ https://genome.jgi.doe.gov/portal/Hosspeendopfungi/Hosspeendopfungi.info.html.

RNA expression values at 3, 6, and 10 days post-inoculation (dpi) were extracted and normalized as described in the original dataset. Transcript per Million (TPMs) values per gene were transformed into Z-scores to compare the different conditions. Heat maps were generated in R using the ComplexHeatmap package ^38^, which performs hierarchical clustering based on Euclidean distance. To identify additional functional domains, the amino acid sequences of the selected proteins were analyzed using PfamScan ^39^, HMMER hmmscan (https://www.ebi.ac.uk/Tools/hmmer/search/hmmscan), and PANTHER ^40^, and the resulting domain architectures were visualized using the drawProteins R package.

### Identification of signal peptides, intrinsically disordered regions, and PrLDs

Signal peptides were predicted using SignalP 6.0 (https://services.healthtech.dtu.dk/services/SignalP-6.0/) ^41^. PrLDs were identified in selected *A. thaliana* and *S. indica* protein sequences using the PLAAC algorithm (http://plaac.wi.mit.edu) ^26^. The minimum core length for prion-like domains was set to 60 amino acids, and the α parameter was set to 50. Intrinsically disordered regions (IDRs) were predicted using IUPred3 (https://iupred.elte.hu), applying default parameters ^42^.

### Gene exon mapping of selected effectors

Raw reads were trimmed with Fastp and trimmed reads were aligned to the *S. indica* genome. The mapped reads were visualized with the Integrative Genomics Viewer (IGV). The Sashimi plot was displayed in IGV by setting the minimum junction depth to 4.

### DNA construct generation

DNA constructs for agroinfiltration assays, stable transgenic plant production, and recombinant protein expression were generated using the Gateway, Golden Gate, or Gibson Assembly cloning systems as indicated. For plant expression, entry clones were generated using the Gateway BP Clonase II Enzyme Mix and recombined into destination vectors using the Gateway LR Clonase II Enzyme Mix (Thermo Fisher Scientific). PEPSI genes were chemically synthesized in pTwist entry vectors (Twist Bioscience), codon-optimized for expression in plant cells, and recombined into pGWB505 or other Gateway-compatible destination vectors depending on the desired C-terminal tag ^43^.

For plant expression of cytosolic GFP, the GFP coding sequence including a stop codon was amplified from pGWB505, subcloned into the Gateway entry vector pDONR221, and subsequently transferred into pGWB502 by LR recombination.

For generating PEPSI12ΔP construct, the coding sequence retaining the native signal peptide but lacking the predicted prion-like domain was amplified by PCR from pENTR_Twist:PEPSI12. The truncated fragment was assembled into the bacterial expression vector pQE80L using Gibson Assembly. During amplification, Gateway recombination sites were incorporated to allow direct transfer of the ΔPrLD fragment into pGWB505 vector.

Plastid-targeted fluorescent control constructs were generated using synthetic genes encoding the HSP21 plastid transit peptide fused to GFP or tdTomato, provided in pTwist entry vectors and transferred into pGWB502 or pGWB402 by LR recombination. Additional synthetic constructs included pENTR_Twist:SP_Pepsi6-mRFP, pENTR_Twist:SP-CFP-PR1-mCherry, and pENTR_Twist:SP-CFP-PEPSI6_CAP-mCherry, were transferred into pGWB502 or pGWB402 as indicated.

For recombinant protein expression, the sequence encoding PEPSI6 lacking the signal peptide was cloned into pPGN-C ^44^, introducing an N-terminal 6×His tag and a C-terminal GST tag. The PEPSI6–GST fragment was amplified by PCR from pGWB523:PEPSI6-GST and assembled into the modified pPGN vector using Gibson Assembly ^45^. The construct was sequence-verified followed by transformation into *E. coli* BL21 for protein expression.

All constructs were sequence-verified and propagated in *Escherichia coli* Mach1 or TOP10 by heat shock prior to downstream applications. A complete list of constructs used in this study is provided in **Supplementary Table 3**.

### Plant material and growth conditions

*Arabidopsis thaliana* wild-type (WT) ecotype Columbia-0 (Col-0) and transgenic lines in the Col-0 background were used for all experiments. Seeds were surface sterilized, stratified at 4 °C in the dark for 3 days, and germinated on half-strength Murashige and Skoog (½ MS) medium containing vitamins (Duchefa Biochemie), adjusted to pH 5.7 and solidified with phytoagar (8 g L⁻¹) (Duchefa Biochemie). Plants were grown under long-day conditions (16 h light/8 h dark) at 22 °C/18 °C with a light intensity of 130 µmol m⁻² s⁻¹. Stable transgenic *Arabidopsis thaliana* lines were generated by Agrobacterium tumefaciens-mediated transformation using the floral dip method, as previously described ^46^. We used FIJI (ImageJ) to measure root length in seedlings grown on vertical agar plates. *A. thaliana* lines used in this study are listed in **Supplementary Table 4**.

### Fungal strains and growth conditions

*Serendipita indica* strain DSM11827 (German Collection of Microorganisms and Cell Cultures, Braunschweig, Germany) was used for all fungal experiments. *S. indica* was cultivated on complete medium (CM) containing 2% (w/v) glucose and 1.5% (w/v) agar, as previously described ^47^. Cultures were incubated at 28 °C in the dark for 4 weeks prior to spore preparation.

### Genome resources

Gene and protein identifiers, accession numbers used in this study are provided in **Supplementary Table 5.**

### Fungal inoculation

Colonization experiments were performed by seed inoculation of *A. thaliana* with *S. indica* spores. Under sterile conditions, *S. indica* spores were scraped from CM agar plates using sterile water containing 0.002% (v/v) Tween 20 (Carl Roth). The spore suspension was washed twice with sterile distilled water and adjusted to a final concentration of 5 × 10⁵ spores mL⁻¹. Mock treatments were performed using sterile distilled water. For seed inoculation, surface-sterilized *Arabidopsis* seeds were incubated in the *S. indica* spore suspension (5 × 10⁵ spores mL⁻¹) for 1 h under sterile conditions, then plated on 1/10 plant minimal nutrition medium (PNM) without sucrose. PNM contained 0.5 mM KNO₃, 0.367 mM KH₂PO₄, 0.144 mM K₂HPO₄, 2 mM MgSO₄·H₂O, 0.2 mM Ca(NO₃)₂, 0.25% (v/v) Fe-EDTA (prepared from 0.56% (w/v) FeSO₄·7H₂O and 0.8% (w/v) Na₂EDTA·2H₂O), and 0.428 mM NaCl. The medium was adjusted to pH 5.7, buffered with 10 mM MES, and solidified with 0.4% (w/v) Gelrite (Duchefa Biochemie).

### RNA extraction and intraradical colonization assay

To quantify intraradical colonization by *S. indica*, root tissue was harvested at the indicated days post inoculation (dpi). Roots were extensively washed with sterile distilled water, and residual surface mycelium was carefully removed using tissue paper. Cleaned roots were immediately snap-frozen in liquid nitrogen and ground to a fine powder.

Total RNA was extracted from ground tissue using the Roboklon RNA extraction kit (Roboklon GmbH, Berlin, Germany) according to the manufacturer’s instructions. RNA concentration and integrity were assessed prior to downstream applications. Complementary DNA (cDNA) was synthesized from total RNA using the RevertAid First Strand cDNA Synthesis Kit (Thermo Fisher Scientific). Quantitative real-time PCR (qRT-PCR) was performed using gene-specific primers to quantify fungal biomass relative to plant material (**Supplementary Table 6)**.

### qRT-PCR analysis

qRT-PCR was performed using a CFX Connect Real-Time PCR Detection System (Bio-Rad). Amplification was carried out under the following conditions: an initial denaturation at 95 °C for 3 min, followed by 40 cycles of 95 °C for 15 s, 59 °C for 20 s, and 72 °C for 30 s, and a final melting curve analysis to verify amplicon specificity.

Relative gene expression levels were calculated using the 2⁻ΔΔCt method ^48^. Primer sequences used for qRT-PCR are listed in the **Supplementary Table 6**.

### Interactome analysis

Roots from 14-day-old *A. thaliana* plants expressing pGFP, PEPSI6–GFP, or PEPSI6ΔP–GFP were used for co-immunoprecipitation. To simulate fungal contact, roots were pre-treated with 25 µM hexaacetyl-chitohexaose (Megazyme) for 1 h prior to protein extraction. Root tissue was harvested and lysed in Co-IP lysis buffer containing 1% Triton X-100, 50 mM Tris-HCl (pH 8.0), 150 mM NaCl, 1 mM EDTA, 1 mM PMSF, and 1X plant protease inhibitor cocktail (Sigma). Lysates were vortexed, clarified by centrifugation at 13,000 × g for 10 min at 4 °C, and supernatants were collected. Equal amounts of total protein from each sample were incubated for 1.5 h with either anti–GFP antibody (1:500; Amsbio) or a control anti-IgG antibody (1:500; Abcam, catalog no. ab46540). Immune complexes were captured by incubation with 50 µL uMACS Protein A MicroBeads (Miltenyi Biotec) for 1 h at 4 °C, then loaded onto pre-cleared µMACS columns (Miltenyi Biotec, catalog no. 130-042-701). Columns were washed three times with wash buffer 1 (50 mM Tris-HCl, pH 7.4, 150 mM NaCl, 5% glycerol, and 0.05% Triton X-100), followed by five washes with wash buffer 2 (50 mM Tris-HCl, pH 7.4, and 150 mM NaCl). Bound proteins were subjected to on-column tryptic digestion using digestion buffer containing 7.5 mM ammonium bicarbonate, 2 M urea, 1 mM DTT, and 5 ng mL⁻¹ trypsin. Peptides were eluted with elution buffer two times (2 M urea, 7.5 mM ammonium bicarbonate, and 15 mM chloroacetamide) and incubated overnight at room temperature with shaking in the dark. The following day, peptides were desalted using StageTips and prepared for label-free quantitative proteomics. For sample data acquisition, we used a Q-Exactive Plus (Thermo Fisher Scientific) mass spectrometer coupled to an EASY nLC 1200 UPLC (Thermo Fisher Scientific), following the protocol detailed at https://www.ebi.ac.uk/pride/archive/projects/PXD041001. MS raw data were processed with MaxQuant (v.5.3.8) ^49^ using default settings with label-free quantification (LFQ) enabled. MS2 spectra were searched against the *A. thaliana* UniProt database, including a list of common contaminants. All downstream analyses were carried out on LFQ values with Perseus. Protein groups were filtered for potential contaminants and insecure identifications. The remaining IDs were filtered for data completeness in at least one group and missing values imputed by sigma downshift (0.3 σ width, 1.8 σ downshift).

### Metabolomics

Roots from 21-day-old seedlings grown under short-day conditions in mock-treated controls were harvested and immediately quenched in liquid nitrogen. Five independent pools, each consisting of approximately 150 mg fresh weight of root tissue (derived from around 250 individual seedlings), were generated and processed for metabolite extraction as described in Balcke et al. ^50^, generating an aqueous and an organic extract. The same day of the extraction 5 μl of the aqueous extract containing hydrophilic metabolites were injected into an Acquity UPLC (Waters Inc.) equipped with a Nucleoshell RP18 column (Macherey and Nagel, 150mm x 2 mm x 2.1μm). Central carbon metabolites including MEP pathway intermediates were separated using tributylammonium as ion pairing agent. Solvent A: 10 mM tributylamine (aqueous) acidified with glacial acetic acid to pH 6.2; solvent B acetonitrile. Gradient: 0-2 min: 2% B, 2-18 min 2-36% B, 18-21 min 36-95% B, 21-22.5 min 95% B, 22.51-24 min 2% B. Column flow was 0.4 ml min^-1^ throughout. The column temperature was 40 °C. Scheduled metabolite detection based on multiple reaction monitoring (MRM) was performed in negative mode with electrospray ionization (ESI) on a QTrap 7500+ (AB-Sciex GmbH, Darmstadt, Germany): Ion source gas 1: 60 psi, ion source gas 2: 70 psi, curtain gas: 35 psi, temperature: 600 °C, ion spray voltage floating: and -2200V). MRM transitions of 189 metabolites covering central carbon and energy metabolism were previously signal-optimized and retention times determined ^50^.

### Quantitative proteomic analysis of Arabidopsis roots

For the quantitative proteomic analysis, root material was collected from 20-day-old *A. thaliana* roots expressing pGFP, PEPSI6–GFP, or PEPSI6ΔP–GFP. For each genotype, samples were obtained under mock conditions. Root material was lyophilized overnight, weighed, and aliquoted into 1.5 mL microcentrifuge tubes, with 2.5 mg of lyophilized tissue per sample.

Protein extraction, purification and digestion were performed using the single-pot solid-phase-enhanced sample preparation (SP4) protocol for proteomics experiments described by Johnston et al. ^51^. Briefly, 2.5 mg of lyophilized tissue was dissolved in 400 µl HCl (0.1 M) and incubated at room temperature for 15 min, shaking. Then, 120 µl of the resulting extract was combined with an equal volume of protein buffer (4% sodium dodecyl sulfate, 0.2 M NaOH) and incubated at 60 °C for one hour, shaking. Protein concentrations were quantified using the Pierce™ BCA Protein Assay Kit (Thermo Fisher Scientific).

The remaining extract was combined with an equal volume of SDT buffer (8% sodium dodecyl sulfate, 0.2 M dithiothreitol, 0.2 M Tris pH 7.6) and incubated at 60 °C for 30 min. The volume containing 30 µg of protein was mixed with 7.5 μl iodoacetamide (0.1 M) and incubated for 30 min in the dark. Then, 2 μl dithiothreitol (0.1 M) was added. Glass beads were used to promote the precipitation of denatured proteins using acetonitrile. The protein-bead pellets were washed multiple times with 80% ethanol. Finally, the proteins were digested for 18 h at 37 °C with 0.5 μg trypsin (MS grade, Promega) in ammonium bicarbonate (50 mM). The peptide-containing supernatant was collected in tubes with low peptide-binding properties. The beads were rinsed with 60 μl ammonium bicarbonate (50 mM) to recover any remaining peptides. The eluates were acidified with formic acid and desalted using 50 mg Sep-Pak tC18 columns (Waters). The purified peptides were quantified using Pierce™ Quantitative Peptide Assay Kit (Thermo Scientific) and adjusted to 400 ng · µl^-1^ in 0.1 % formic acid.

For the Quantitative Shotgun Proteomics by Ion Mobility Mass Spectrometry (IMS-MS/MS), 400 ng of peptides were injected via a nanoElute2 UHPLC (Bruker Daltonics) and separated on an analytical reversed-phase C18 column (Aurora Ultimate 25 cm x 75 µm, 1.6 µm, 120 Å; IonOpticks). Using MS grade water and a multi-staged gradient acetonitrile containing 0.1 % formic acid (0 min, 2%; 54 min, 25%; 60 min, 37%; 62 min, 95%; 70 min, 95%). Peptides were eluted and ionized by electrospray ionization with a CaptiveSpray2 ion source and a flow rate of 300 nl · min^-1^. The ionized peptides were separated, fragmented and analyzed with a standard data-dependent acquisition parallel accumulation-serial fragmentation application method (DDA-PASEF) of the system with the following settings: Ion mobility window of 0.7 - 1.5 V · s · cm^-2^, 4 PASEF ramps, target intensity 14,500 (threshold 1,200), and a cycle time of ∼0.53 s. The analysis was performed on a timsTOF-HT mass spectrometer (Bruker Daltonics).

Peptide and protein identification were performed using MaxQuant software (version 2.4.13.0; ^52^. Search parameters were set automatically by loading the raw files (.d) acquired on the timsTOF-HT instrument. Database searching was performed simultaneously against FASTA files from *Arabidopsis thaliana* (TAIR10; Arabidopsis.org), *Serendipita indica* (UP000007148_2024_07_15; UniProt), and the PEPSI6–GFP constructs.

Protein quantification was carried out using label-free quantification (LFQ), and the calculation of intensity-based absolute quantification (iBAQ) values was enabled. The Match Between Runs option was disabled to avoid erroneous peptide transfer between runs originating from different proteomes.

Data evaluation was performed using Perseus software (version 2.1.4.0; ^53^. Protein groups were excluded from further analysis if they were not detected in at least four biological replicates in at least one sample group. Missing values were subsequently imputed by random sampling from a normal distribution. Fold changes between sample groups were calculated based on log₂-transformed LFQ intensities using Student’s t tests with a false discovery rate (FDR) of 0.05.

### Protein expression and purification

A single colony was used to inoculate a 60 mL preculture, which was used to set up two 1 L expression cultures (OD600 = 0.01). Cultures were grown at 37 °C and 200 rpm to OD600 ≈ 0.4, induced with 400 µM IPTG, and incubated for 4 h at 30 °C. Cells were harvested by centrifugation (4,500 rpm, 20 min, 4 °C). Pellets were resuspended in Ni-NTA lysis buffer pH 8.0 (50 mM NaH₂PO₄, 300 mM NaCl, 10 mM imidazole), lysed by sonication (10 min, pulse mode), and clarified by centrifugation (20,000 rpm, 25 min, 4 °C). The supernatant was applied to a 1 mL HisTrap FF Crude column (Cytiva) using an ÄKTA Start system. The column was equilibrated with 5 CV binding buffer pH 7.4 (20 mM Na₃PO₄, 500 mM NaCl, 25 mM imidazole), washed with 15 CV, and eluted isocratically with 100% 5 CV elution buffer pH 7.4 (20 mM Na₃PO₄, 500 mM NaCl, 500 mM imidazole). 1 ml fractions were analyzed by SDS-PAGE. Fractions containing His-PEPSI6-GST (without SP) were pooled, concentrated as required, and further purified by size-exclusion chromatography on a Superdex 75 pg HighLoad column (Cytiva) using an ÄKTA system using SEC running buffer pH 7.5 (150 mM Tris-HCl, 600 mM NaCl). 5 ml fractions were collected and analyzed by SDS-PAGE. Fractions containing His-PEPSI6-GST were pooled and concentrated. Protein concentration was determined by absorbance at 280 nm. Glycerol was added to 10% (v/v) prior to storage at −80 °C.

### Western blotting analysis of plants

Plant material was ground in liquid N_2._ The powder was resuspended in ice-cold TKMES homogenization buffer (100 mM Tricine-potassium hydroxide, pH 7.5, 10 mM KCl, 1 mM MgCl2, 1 mM EDTA and 10% (w/v) sucrose) supplemented with 0.2% (v/v) Triton X-100, 1 mM DTT, 100 µg ml^−1^ of phenylmethylsulfonyl fluoride (PMSF), 3 µg ml^−1^ of E64 and plant protease inhibitor. After centrifugation at 10,000*g* for 10 min (4 °C), supernatant was collected for a second centrifugation. Protein concentration was determined with Pierce Coomassie Plus (Bradford) Protein Assay Kit. Total protein was SDS–PAGE separated, transferred to a PVDF membrane (BioRad) and subjected to western blotting. The following antibodies were used: anti–GFP (AMSBIO, catalog no. TP401, 1:5,000), anti-Actin (Agrisera, catalog no. AS132640, 1:5,000), anti-UGPase (Agrisera, catalog no. AS23 4916, 1:1,000) anti-GST (Sigma, catalog no. G7781, 1:2,000).

### Sequence alignment of CAP proteins

Amino acid sequences of CAP-domain–containing proteins from *Ustilago maydis*, *S. indica* and *A. thaliana* were retrieved from publicly available databases. To focus on conserved regions, predicted signal peptides and intrinsically disordered regions were excluded prior to alignment where indicated. Multiple sequence alignments were performed using Clustal Omega ^54^ with default parameters. Alignments were visualized and manually curated using GeneDoc 2.7 ^55^ and ESPript 3.0 ^56^. Conserved regions and putative CAPE peptide motifs were identified based on sequence conservation and comparison with previously characterized plant ^57^ and fungal ^31^ CAP proteins.

### Statistical analyses

Statistical analyses were performed using GraphPad Prism version 10 (GraphPad Software). Comparisons between two groups were conducted using two-tailed Student’s t tests, as specified in the corresponding figure legends. For comparisons involving more than two groups, ordinary one-way ANOVA was used; when a significant effect was detected, Tukey’s multiple-comparisons test was applied for post hoc analysis. Statistical differences identified by Tukey’s test are indicated by different letters in the figures. Sample sizes (*n*) and exact *P* values are reported in the figures or figure legends.

### Microscope setup and live-imaging

For 3D live imaging, confocal microscopy was performed using an inverted Leica STELLARIS microscope equipped with a Plan-Apochromat 63X/1.4 NA oil-immersion CS2 objective and a galvano stage (Fig. 4f). Prior to analysis, Leica FALCON (Fast FLIM) phasor separation was applied to the autofluorescence channel to enhance signal discrimination.

Additional 3D live-imaging experiments were carried out on an inverted Zeiss LSM 880 Airyscan microscope using a Plan-Apochromat 63X/1.4 NA oil-immersion DIC M27 objective and a piezo stage (Fig. 4c). Spectral imaging was performed using the LSM 880 GUI spectral unmixing module to separate emission spectra from background signals, followed by Airyscan deconvolution to improve spatial resolution.

For 4D live-imaging time-lapse experiments, image acquisition was performed using an inverted Nikon AX-R NSOARC microscope equipped with a 60×/1.27 NA oil-immersion objective and a piezo stage, and operated in resonance scanning mode with 8X averaging (unidirectional) and denoising (**Supplementary Video 1**). The video shows merged z-stacks.

All images were processed and exported as TIFF files using ImageJ (NIH) prior to analysis. Brightness and contrast were adjusted equally across entire images. All image analyses were performed using ImageJ (NIH).

For all 3D and 4D live imaging, roots were mounted under 95% perfluorodecalin (Sigma-Aldrich, P9900-25G) in a µ-Slide 1 Well Glass Bottom #1.5H (170 µm ± 5 µm), D 263 M Schott glass (ibidi, Art. No. 82107), and covered with a piece of Lumox Film 25 (Sarstedt AG; Ref. 94.6077.317).

Fluorescence intensity profiles of eGFP and chlorophyll autofluorescence were extracted along defined line scans across plastids or regions of interest. Fluorescence intensity values were plotted as a function of distance (µm) to assess signal distribution and overlap. Quantification of fluorescence intensity was performed using Leica Application Suite X (LAS X) software (version 3.7.4.23463).

For colocalization analyses, orthogonal (x–z and y–z) views were generated from confocal *z*-stacks using Leica LAS X software to confirm spatial overlap of fluorescent signals within plastids. Imaging parameters were adjusted as needed to accommodate differences in fluorophore expression levels between samples. Image analysis was performed on raw data without additional normalization unless otherwise stated.

The following fluorophores were detected using appropriate excitation and emission settings: eGFP, eCFP, mRFP, mCherry, and chlorophyll autofluorescence. For visualization of fungal structures, *S. indica* hyphae were stained with wheat germ agglutinin conjugated to Alexa Fluor 555 (WGA–AF555, ThermoFisher Scientific).

### Structural visualization and analysis of PEPSI6 and PR1

Predicted protein structures of *Serendipita indica* PEPSI6 and *A. thaliana* PR1 were obtained from AlphaFold3 server ^58^. Structural visualization and analysis were performed using PyMOL (Schrödinger, LLC). Signal peptides were excluded from the structural models prior to visualization. Structural features, including the CAP domain, were highlighted and compared between PEPSI6 and PR1. Structural representations were generated using standard PyMOL rendering modes, and figures were prepared for publication using PyMOL.

### Activity-based protein profiling (ABPP) and *in vitro* cleavage assays

Arabidopsis RD21C and a catalytically inactive RD21A mutant (RD21A^mut^) were transiently expressed in *N. benthamiana* by Agrobacterium-mediated transformation. Leaves of 4–6-week-old plants were infiltrated on the abaxial side with bacterial suspensions (OD600 = 1) using a needleless syringe. Leaf material was harvested 3 days post infiltration. Apoplastic fluid was isolated by vacuum infiltration of leaves in water (200 mbar, 10 min, three cycles), followed by centrifugation at 2000 × g for 15 min using syringe-based collection.

For ABPP and *in vitro* cleavage assays, equal volumes of apoplastic fluid were mixed with NaOAc buffer (pH 6.0; final concentration 50 mM) and DTT (final concentration 10 mM). Samples were preincubated in the presence or absence of 30 µM E64 for 20 min followed by the addition of MV201 ^59^ to a final concentration of 2 µM. Samples were labeled for 2 h at room temperature in the dark. Reactions were terminated by the addition of 1X Laemmli buffer. Proteins were resolved by SDS–PAGE, followed by fluorescence scanning using a Bio-Rad ChemiDoc system and Sypro® Ruby staining.

For *in vitro* cleavage assays, 1.25 µg recombinant His–PEPSI6–GST (without SP) was incubated for 0, 15, or 30 min at room temperature. Reactions were stopped with 6× Laemmli buffer, and samples were analyzed by SDS–PAGE and Western Blot using a GST antibody. Membranes were stained by Coomassie brilliant blue.

## Data Availability

All data supporting the findings of this study are available within the paper and its Supplementary Information files. Mass spectrometry–based proteomics data generated in this study have been deposited in the ProteomeXchange Consortium via the PRIDE partner repository. Two independent datasets are available: (i) co-immunoprecipitation–based interactomics of GFP, PEPSI6–GFP and PEPSI6ΔP–GFP complexes in *A. thaliana* roots (accession number PXD074150) and (ii) global label-free quantitative proteomics of *A. thaliana* roots expressing pGFP, PEPSI6–GFP, and PEPSI6ΔP–GFP (accession number PXD072775).

## Author contributions

E.L. conceived and designed the project, performed most experiments including microscopy, analyzed and visualized the data, wrote the manuscript, and supervised the work. T.K. performed imaging experiments, generated and characterized transgenic lines, cloned the ΔPrLD constructs, performed root measurements, and conducted colonization assays. M.K. and J.M.V. cloned and purified PEPSI6–GST, performed ABPP and *in vitro* cleavage assays. C.A., B.H., and T.M.H. performed and analyzed the total proteomics experiments. S.K. and D.V. contributed to the design and optimization of the co-immunoprecipitation experiments. J.K. performed imaging experiments, colonization assays and western blot analyses. D.M. cloned effector constructs, conducted agroinfiltration experiments, and generated confocal microscopy data. P.K. performed confocal microscopy of PEPSI6–mRFP and DXR–GFP double transgenic lines. T.N. analyzed transcriptomic data, generated heat maps and performed structural analyses. G.U.B. performed metabolomic analyses. G.L. performed sequence alignments. A.Z. conceived and supervised the project, contributed to experimental design and interpretation, secured funding, and co-wrote the manuscript. All authors discussed the results and contributed to the final manuscript.

## Acknowledgments

E.L., M.K., C.A., P.K., B.H., T.M.H., J.M.V., and A.Z. acknowledge funding from the Deutsche Forschungsgemeinschaft (DFG, German Research Foundation) under Germany’s Excellence Strategy – EXC-2048/2 – project ID 390686111. The proteomics unit in T.M.H.’s laboratory is funded by DFG-INST 216/1290-1 FUGG. E.L. is supported by the German Federal Ministry of Education and Research (BMBF) through the GO-Bio initial funding program (grant 16LW0639). A.Z. acknowledges support from iHEAD (NRW Profilbildung ID: PB22-025A). The funders had no role in study design, data collection and analysis, decision to publish, or preparation of the manuscript.

We thank Manuel Rodríguez-Concepción for providing the DXR–GFP *Arabidopsis* lines; Renier van der Hoorn for providing the RD21 constructs; Herman Overkleeft for providing MV201; and Tsuyoshi Nakagawa for providing the pGWB vectors. We thank Concetta De Quattro for help with sashimi plot generation; Sinaeda Anderssen for support with bioinformatic analyses, prion-like domain predictions, and protein domain visualizations; Ashmita Ray for technical support; Nyasha Charura and Tim Thomsen for helpful discussions; and Gunther Döhlemann for facilitating laboratory space and materials.

We thank the CEPLAS Biocenter Imaging Facility for technical support and imaging services. We also thank Prerana Wagle, Jan-Wilm Lackman, and Stefan Müller from the CECAD/CMMC Proteomics Facility for support, analysis, and advice on proteomics experiments. The proteomics facility was supported by the DFG large-instrument grant INST 216/1163-1 FUGG.

During preparation of this manuscript, ChatGPT (OpenAI) was used to assist with language editing and readability. All content was reviewed and edited by the authors, who take full responsibility for the final manuscript.

