## Supplemental Figures 1-11 for "Prion-like domains control plastid targeting of PEPSI effectors and PEPSI6-mediated modulation of DXR during root symbiosis"

**PIIN_01066 - PEPSI4 - *A. thaliana* with *S. indica* 10 dpi**

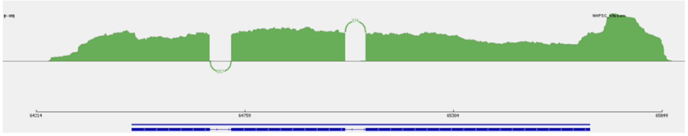


MTLFNSLLSAALLAVSVLTAQADATPLVANGLSARHPHVGMSANHHGQLRKRVSHRHRKRCLAQNAPGASSSSSSAPPAPTGNSNDGNNNNNNNNNNGGNNNNGGGNTYTPPPIGGAVCGDKKLGLAAEWNVDAS-
RLVSPRTCITYNWDASNRGPYDIPGLIYVPMVSRLGNLDGFYANVVNRPDRNYPWIAFLNEANEPGQANVDVGTAVALWRQHALPLRNRGIKLLSPSVTSSPEGTRWIVDFYNSLGDWEKPDAISLHWYGTDFDSL-
RNYINDMSGRFGGRHIWLTEFGCTLFPYVPCSIDQGSFAAQAIAFLNSHWAIDIYAPFGFVNDLGGANEGVRLIDPGSGNANSLGSMYLWS

**PIIN_09841 - PEPSI6 - *A. thaliana* with *S. indica* 10 dpi**

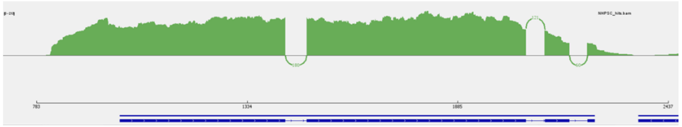


MIALKSTVVVLSLLASQATAVPTFDAASPASNEARGLVDDATNAIQRVWEGIKCLGDCRKSWGWTGNHFGGDPWGPPLKIGDPVPYTPSEDSSGIVSTTTTARSTTRTSSAEGGFGILNVDTSRTTLVDGSN-
VEFSVFNVPTLTRSTSTSTIVLPTSTTPPPPPPEPTTTFQEPQPSPEPQPSSSQEQPQPQEQPAPSPTPSNNDNNNNNNVGNNNSGGSTQGAAGDIAAYLAAHNSARSQHGASPVSWSDELAGFAQEWANNC-
QFQHSQGKFGRVGENLAAGTGQYSIEDMVGDWVAEVTDYNPSNPKASHFTQVVWKATTQIGCAKQTCTGIFGNTPATYYVCEYREAGNVIGNFAANVQA

**PIIN_05452 - PEPSI12 - *A. thaliana* with *S. indica* 10 dpi**

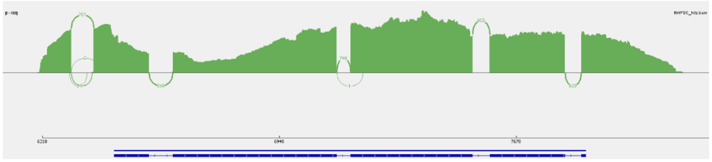


MKLSLSLATVLTLLPMLVTASSSPFIERSIHDTAYLPRVARRFGRNPDIFGRQNNAADRAATAQRAAQQAAAQRAQQEQQAAIDRQRAAIASSAAAQQSAAAERNRQADIANGNNRNGGNGGNQQNNNGQNNNNNN-
GATRTTSAAAAATTTANNGGNNNGNNNNNNGGNNNNGGNNNNGGNNNNGGNNGGNNGGNNGGNNNNAITLDPSLVASNLAKTGQETPAAGQVASLTSTNNFLNFCAGKTITNGQQITTGSCNPIPIGDVAPKANMP-
SCKFVNPINFSTIPANRDFTIQMKLNNLQAGVFTNAQKTYYMAPQQLNAQGDIIGHTHFVVQKMGAFDDPTILDAGTFQFFKGINTAAVNGIVSTTVTGGIPPGNYRLASINTDANHAPAIVAVAQHGFVDDMIYF-
TAE

**Extended Data Fig. 1. Transcript structure and splicing of selected PEPSI genes during plant interaction.** Sashimi plots showing RNA-seq read coverage and splice junction usage across exons and introns of selected *Serendipita indica* PEPSI genes (PEPSI4, PEPSI6, and PEPSI12). Data were obtained from *S. indica* grown in contact with *Arabidopsis thaliana* roots at 10 days post inoculation (dpi). Green tracks indicate read coverage across genomic loci, and arcs represent splice junctions, with thickness proportional to the number of supporting reads. Raw reads were trimmed using Fastp and aligned to the *S. indica* reference genome. Aligned reads were visualized in the Integrative Genomics Viewer (IGV), and sashimi plots were generated with a minimum junction read depth of 4.

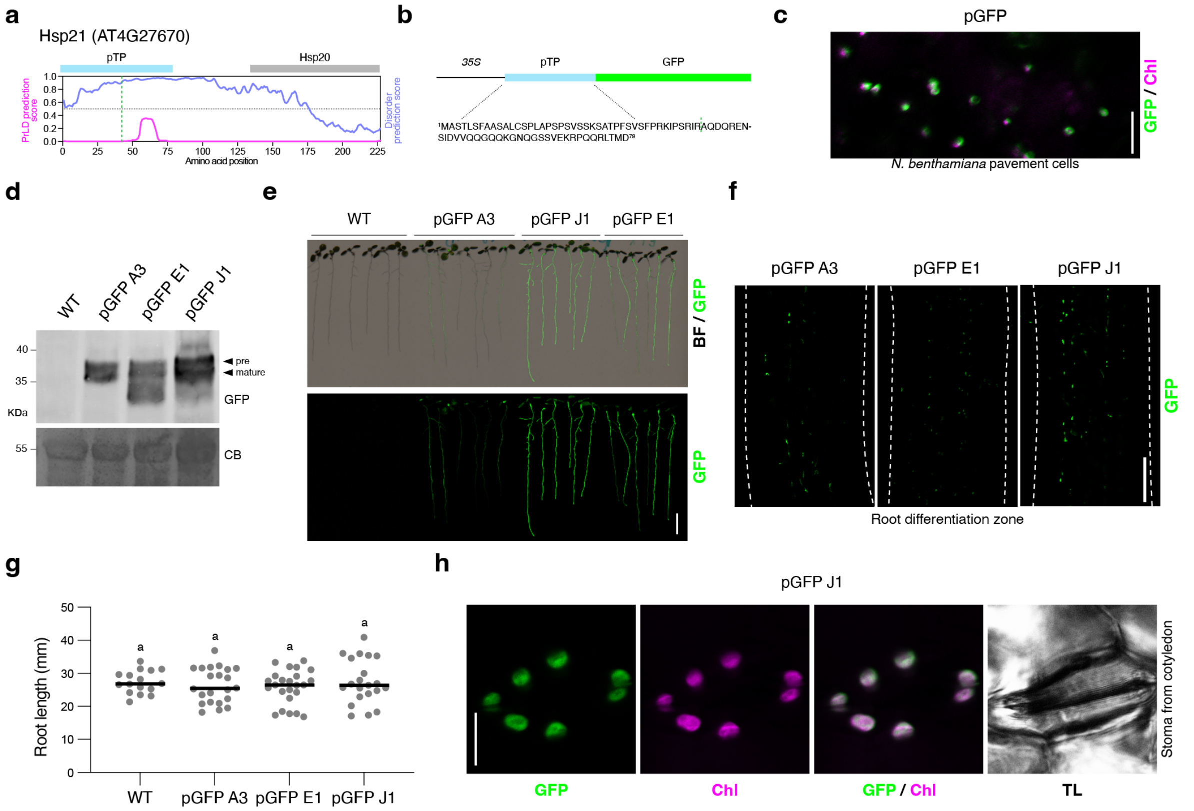


**Extended Data Fig. 2. Characterization of plastid-targeted GFP control lines. a**, Prediction profiles for prion-like domains (PrLDs) and intrinsically disordered regions (IDRs) of the chloroplast heat shock protein Hsp21 (AT4G27670). The predicted plastid transit peptide (pTP) is shown in blue, and the Hsp20 domain is indicated in gray. **b**, Schematic representation of the pGFP construct showing the predicted plastid transit peptide (pTP) fused to GFP under the control of the 35S promoter. The amino acid sequence of the transit peptide is shown below, and the predicted cleavage site is indicated by a blue dashed line. **c**, Representative confocal microscopy image of *Nicotiana benthamiana* leaf pavement cells transiently expressing pGFP, acquired 3 days after agroinfiltration. GFP fluorescence (green) and chloroplast autofluorescence (magenta) are shown. Scale bar, 20 µm. **d**, Immunoblot analysis of total protein extracts from wild-type (WT) plants and independent pGFP transgenic lines using an anti-GFP antibody. Coomassie Brilliant Blue staining showing the Rubisco large subunit (RbcL) is included as a loading control. Representative of two independent experiments. **e**, Representative images showing GFP fluorescence in roots of 10-day-old independent pGFP transgenic lines. Images were acquired using a LI-COR Odyssey M imager in bright-field (BF) and Alexa Fluor 488 (green) channels. Scale bar, 5 mm. **f**, Confocal microscopy images of the differentiation zone of 10-day-old pGFP roots. GFP fluorescence is shown in green. Scale bar, 50 µm. **g**, Root length analysis of 10-day-old WT and pGFP lines. The scatter plot shows individual measurements with the median indicated. Statistical analysis was performed using ordinary one-way ANOVA to compare group means. No statistically significant differences were detected ($F(3,79) = 0.4541$, $P = 0.7152$; WT n = 16, pGFP A3 n = 22, pGFP E1 n = 24, pGFP J1 n = 21). **h**, Representative confocal microscopy images of stomata from cotyledons of 10-day-old pGFP seedlings. GFP fluorescence (green), chloroplast autofluorescence (magenta), merged images, and transmitted light (TL)

are shown. Scale bar, 10 µm. All images are representative of three independent experiments unless stated otherwise.

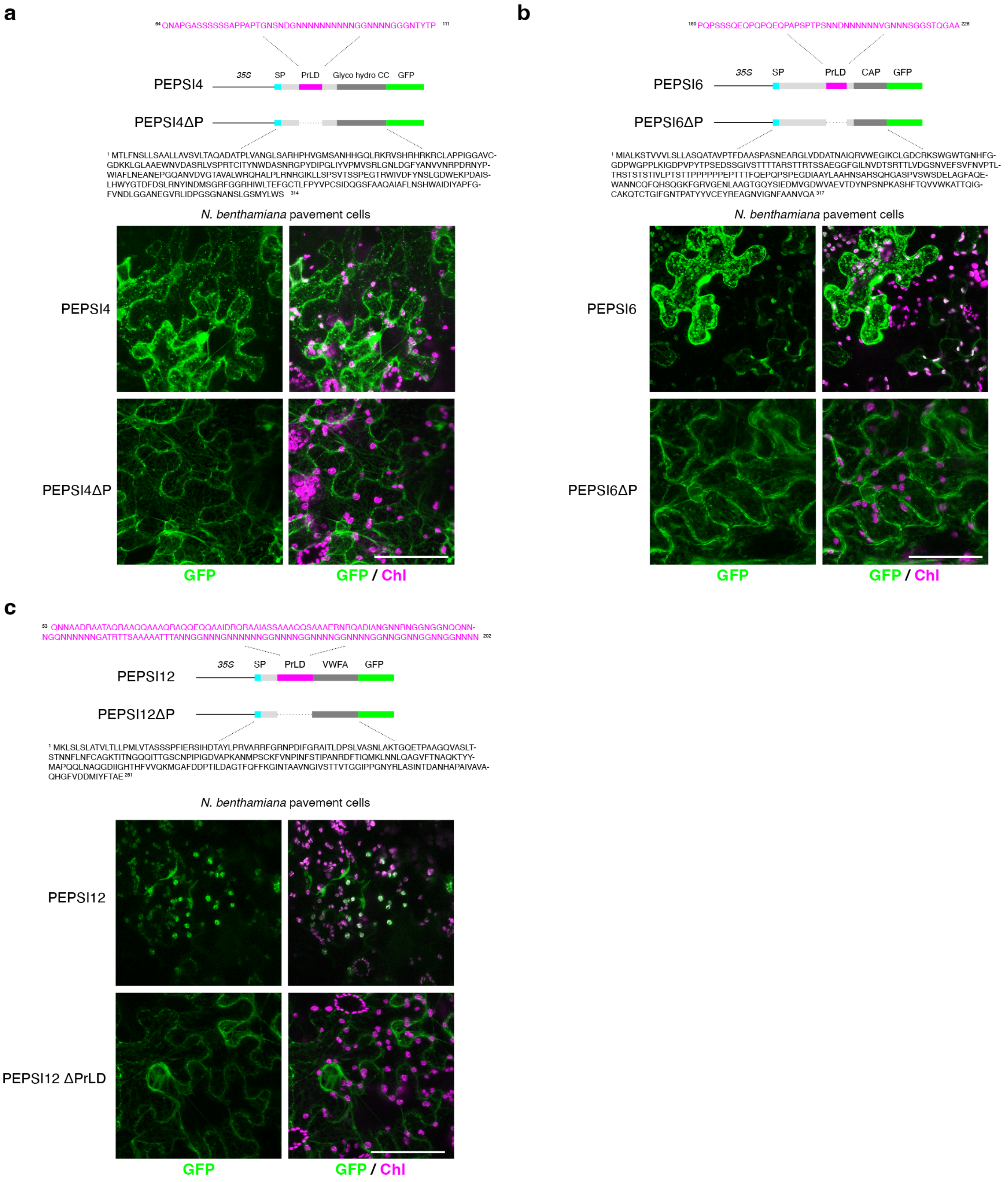


**Extended Data Fig. 3. Prion-like domains are required for plastid localization of PEPSI effectors in *Nicotiana benthamiana*.** Representative confocal microscopy images of *N. benthamiana* epidermal pavement cells transiently expressing GFP-tagged PEPSI effectors and their prion-like-domain deletion variants. **a**, PEPSI4–GFP and PEPSI4ΔPrLD–GFP. **b**, PEPSI6–GFP and PEPSI6ΔPrLD–GFP. **c**, PEPSI12–GFP and PEPSI12ΔPrLD–GFP. Images show GFP fluorescence alone and merged with chlorophyll autofluorescence (Chl). Scale bars, 50 µm. Images are representative of three independent experiments. For each panel, schematic representations of the corresponding constructs are shown above the images. Cyan boxes indicate signal peptides (SP), magenta boxes denote prion-like domains (PrLD), dark gray boxes represent additional predicted functional domains, and green boxes represent GFP. PrLD amino acid sequences are shown in magenta, and the sequences of the corresponding PrLD-deletion constructs are displayed below.

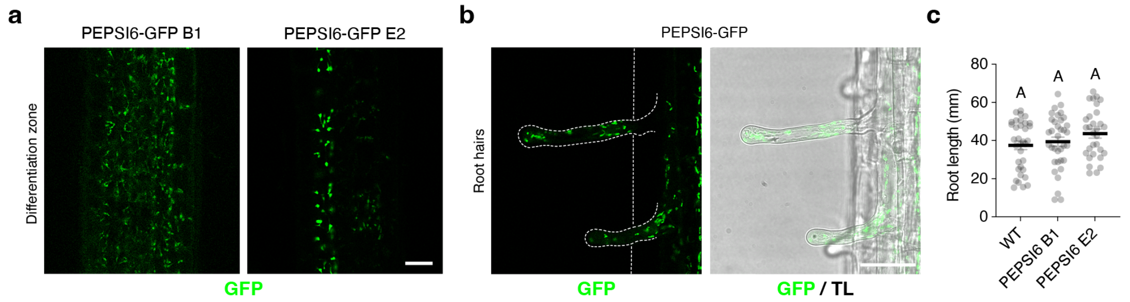


**Extended Data Fig. 4. Characterization of PEPSI6–GFP transgenic Arabidopsis lines.** **a**, Representative confocal microscopy images showing PEPSI6–GFP fluorescence in the differentiation zone of roots from 7-day-old independent transgenic lines (B1 and E2). Images show the GFP channel (green). Scale bar, 20 µm. **b**, Representative images showing PEPSI6–GFP localization in root hairs. GFP fluorescence (green) and merged GFP with transmitted light (TL) are shown. Scale bar, 50 µm. **c**, Root length analysis of 12-day-old wild-type (WT) and PEPSI6–GFP lines grown under long-day (LD) conditions. The scatter plot shows individual measurements with the median indicated. Statistical analysis was performed using ordinary one-way ANOVA to compare group means. No statistically significant differences were detected ($F(2,96) = 1.769$, $P = 0.1760$; WT $n = 34$, PEPSI6 B1 $n = 35$, PEPSI6 E2 $n = 30$).

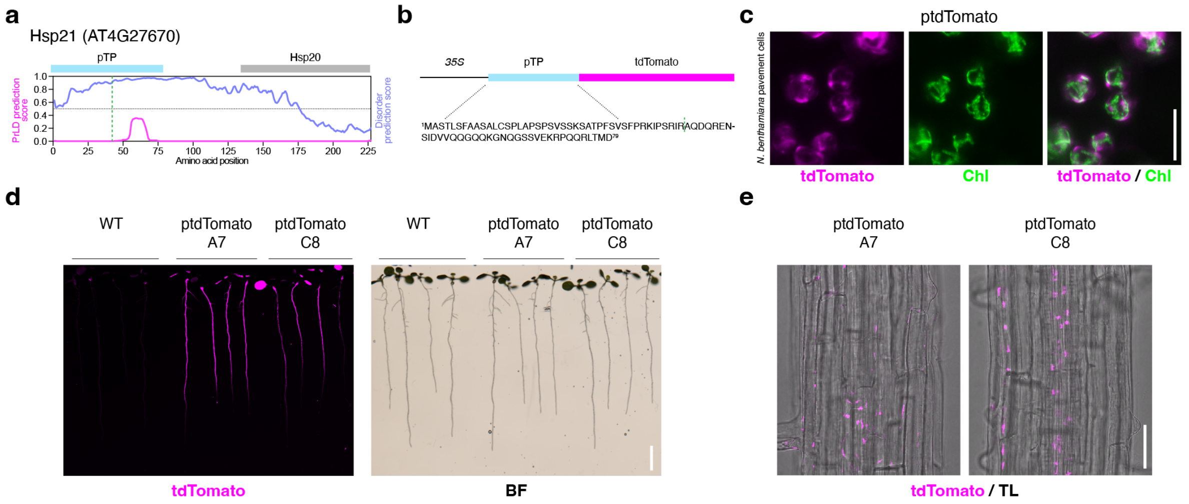


**Extended Data Fig. 5. Characterization of a plastid-targeted tdTomato reporter line.** **a**, Prediction profiles for prion-like domains (PrLDs) and intrinsically disordered regions (IDRs) of the chloroplast heat shock protein Hsp21 (AT4G27670). The predicted plastid transit peptide (pTP) is indicated in blue, and the Hsp20 domain is shown in gray. **b**, Schematic representation of the plastid-targeted tdTomato (ptdTomato) construct, showing the Hsp21-derived chloroplast transit peptide fused to tdTomato under the control of the 35S promoter. The amino acid sequence of the transit peptide is shown below, and the predicted cleavage site is indicated by a blue dashed line. **c**, Representative confocal microscopy images of *Nicotiana benthamiana* leaf pavement cells transiently expressing ptdTomato, acquired 3 days after agroinfiltration. tdTomato fluorescence is shown in magenta, chlorophyll autofluorescence in green, and merged images are shown on the right. Scale bar, 10 µm. **d**, Generation of Arabidopsis plastid ptdTomato transgenic lines. Representative images show tdTomato fluorescence (left) and corresponding bright-field (BF) images (right) of 10-day-old wild-type (WT) seedlings and two independent ptdTomato lines (A7 and C8). Scale bar, 5 mm. **e**, Confocal microscopy images of roots from 10-day-old ptdTomato transgenic lines (A7 and C8), showing tdTomato fluorescence (magenta) overlaid with transmitted light (TL). Scale bar, 50 µm. All images are representative of at least three independent experiments.

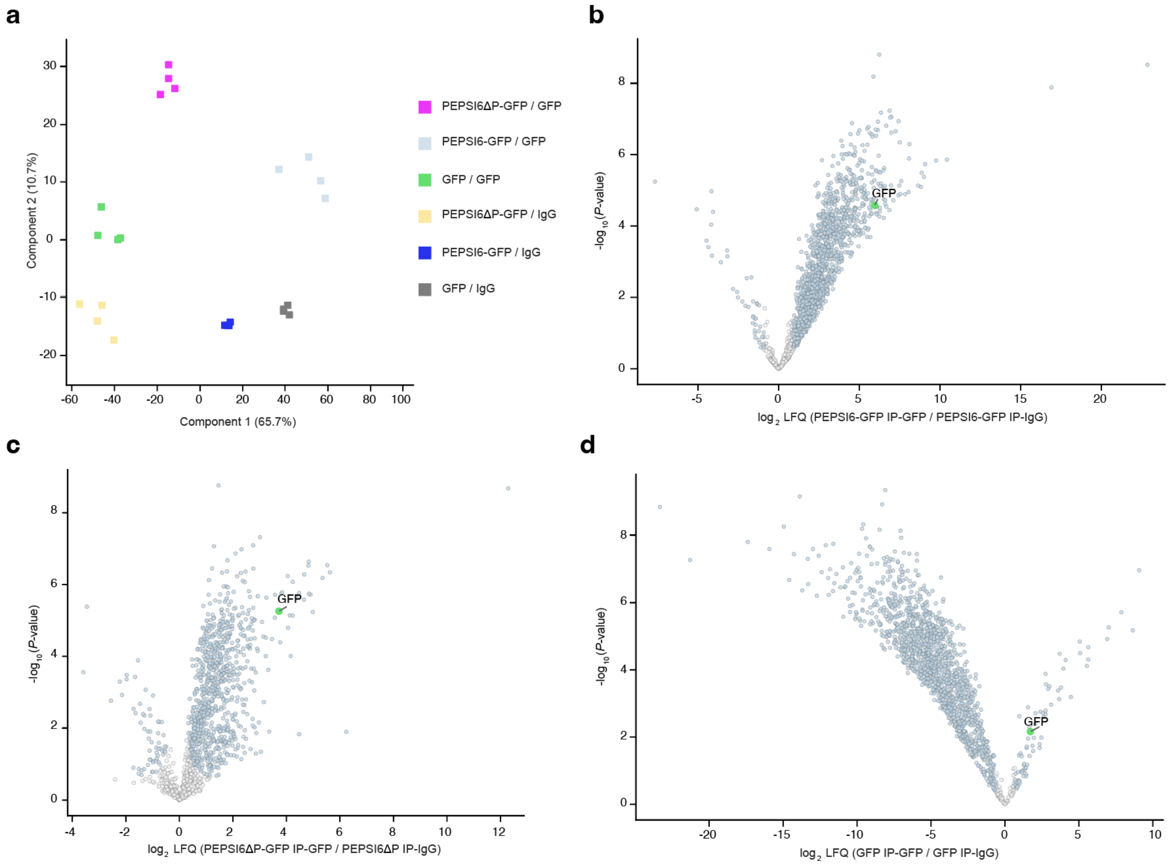


**Extended Data Fig. 6. Validation of PEPSI6–GFP and PEPSI6ΔP-GFP protein interactions by co-immunoprecipitation. a**, Principal component analysis (PCA) of label-free quantitative (LFQ) proteomics data obtained from co-immunoprecipitation (co-IP) experiments using an anti-GFP antibody. Protein complexes were isolated from roots of Arabidopsis lines expressing PEPSI6-GFP, PEPSI6ΔP-GFP, or cytosolic GFP, with IgG pulldowns serving as negative controls. Seven-day-old roots were pretreated with chitohexaose (25 µM) for 1 h prior to protein extraction. Distinct clustering of PEPSI6–GFP, PEPSI6ΔP-GFP, and GFP samples is observed (n = 4 biological replicates). **b–d**, Volcano plots comparing specific anti-GFP pulldowns with corresponding IgG controls for (**b**) PEPSI6-GFP, (**c**) PEPSI6ΔP-GFP, and (**d**) cytosolic GFP samples. Plots show $-\log_{10}$(P value) versus $\log_2$ fold enrichment of protein abundance based on LFQ intensities. Statistical significance was assessed using a two-tailed Student's t test with four biological replicates, and proteins with an FDR-adjusted q value < 0.05 were considered significantly enriched. GFP is highlighted (green dot) among the significantly enriched proteins (blue dots), validating the specificity of the immunoprecipitation.

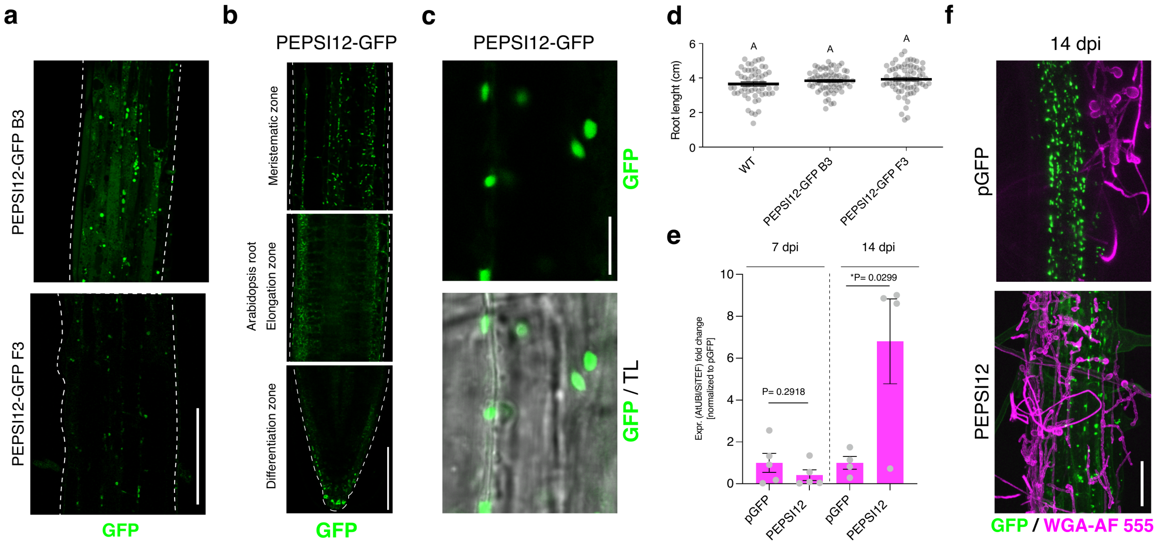


**Extended Data Fig. 7. Characterization of PEPSI12 transgenic Arabidopsis lines.** **a,** Confocal microscopy images of roots from two independent *Arabidopsis thaliana* transgenic lines, B3 and F3, expressing PEPSI12–GFP. GFP signal is shown in green. Scale bar, 100 µm. **b,** Representative confocal microscopy images of 7-day-old *A. thaliana* roots expressing PEPSI12–GFP, showing the meristematic, elongation and differentiation zones of the root. Scale bar, 100 µm. **c,** Magnified views of cells in the differentiation zone of roots expressing PEPSI12–GFP. GFP fluorescence is shown alone and merged with transmitted light (TL). Scale bar, 10 µm. **d,** Root length analysis of independent PEPSI12–GFP transgenic lines. Points represent individual roots and bars indicate the median. Statistical significance was assessed by one-way ANOVA followed by Tukey's multiple-comparisons test. **e,** Intraradical *Serendipita indica* colonization quantified by qPCR in seed-inoculated pGFP control and PEPSI12–GFP plants at 7 and 14 days post inoculation (dpi). Values were normalized to pGFP control plants. Statistical significance was assessed using a two-tailed unpaired Student's *t*-test. **f,** Representative confocal microscopy images of *S. indica*-colonized roots. GFP fluorescence is shown in green and fungal structures are shown in magenta. Scale bar, 50 µm.

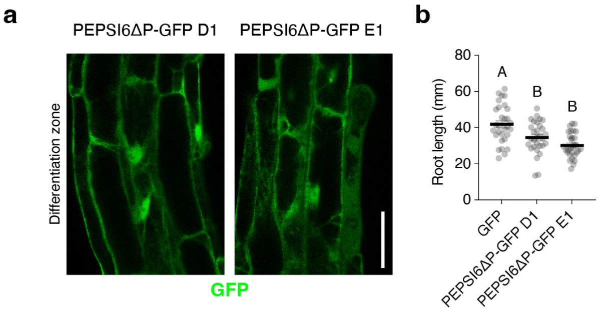


**Extended Data Fig. 8. Characterization of PEPSI6ΔP–GFP transgenic Arabidopsis lines.** **a**, Representative confocal microscopy images showing PEPSI6ΔP–GFP fluorescence in the differentiation zone of roots from 7-day-old independent transgenic lines (D1 and E1). Images show the GFP channel (green). Scale bar, 50 µm. **b**, Root length analysis of 12-day-old GFP control and PEPSI6ΔP–GFP lines grown under long-day (LD) conditions. Root length was compared across genotypes using ordinary one-way ANOVA followed by Tukey's multiple-comparisons test ($F(2,87) = 13.45$, $P < 0.0001$; n = 30 seedlings per genotype). Different letters indicate statistically significant differences between groups.

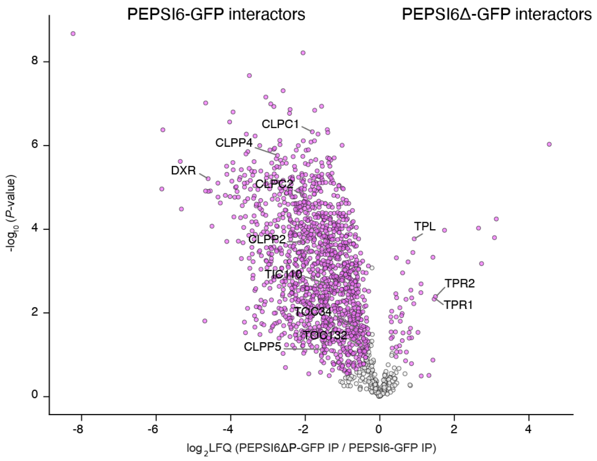


**Extended Data Fig. 9. Comparison of PEPSI6–GFP and PEPSI6ΔP–GFP protein interactomes.** Volcano plot comparing proteins co-immunoprecipitated with PEPSI6–GFP and PEPSI6ΔP–GFP from 7-day-old Arabidopsis roots. The plot shows $-\log_{10}$(P value) from a two-tailed Student's t test plotted against the $\log_2$ fold change of protein abundance based on label-free quantification (LFQ) intensities, calculated as PEPSI6ΔP–GFP relative to PEPSI6–GFP. Magenta dots indicate proteins showing significant differences in enrichment between the two samples (Student's t test; n = 4–5 biological replicates).

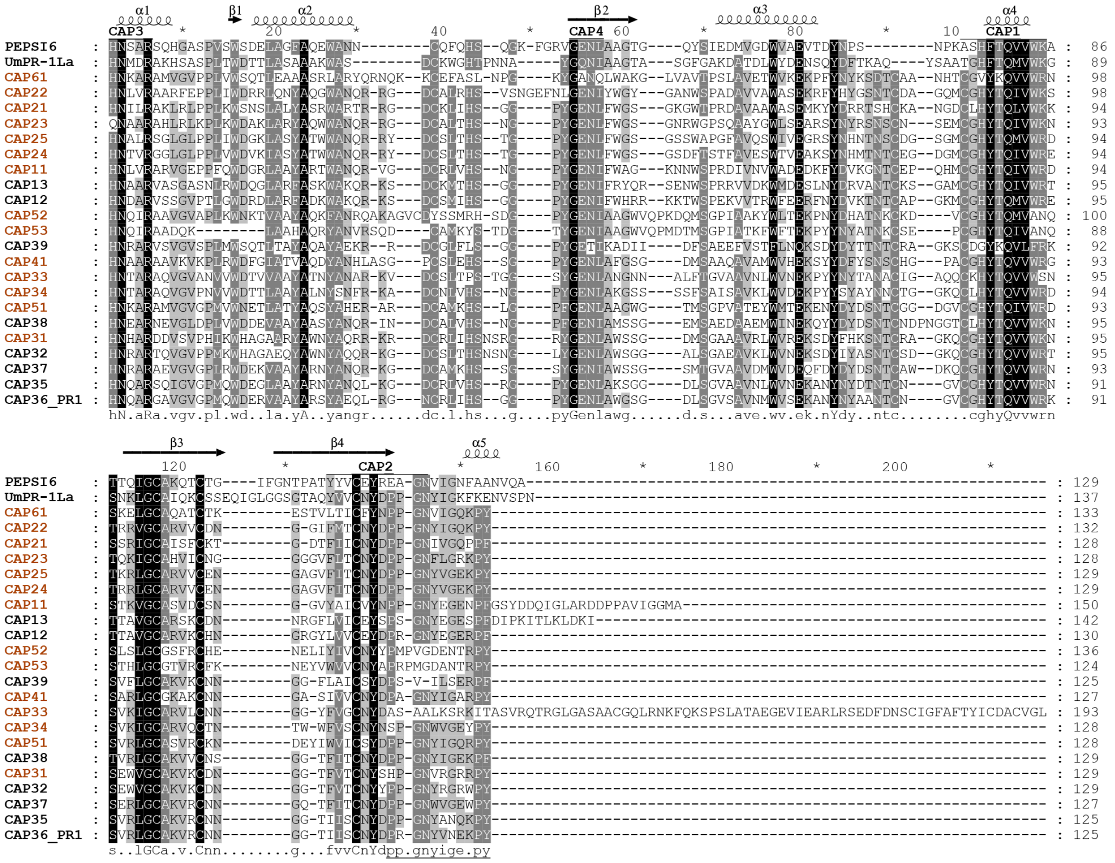


**Extended Data Fig. 10. Sequence alignment of CAP proteins from plants and fungi**. Multiple sequence alignment of selected CAP proteins from *A. thaliana* (Matsuzawa *et al.*, 2024)*, S. indica PEPSI6, and Ustilago maydis UmPR-1La* (Lin *et al.*, 2023). Predicted secondary structure elements (α-helices and β-sheets) are indicated for PEPSI6 above the alignment. Arabidopsis CAP proteins classified as root expressed by Matsuzawa et al. (2024) are highlighted in brown. The alignment was performed after removal of signal peptides and intrinsically disordered regions (IDRs).

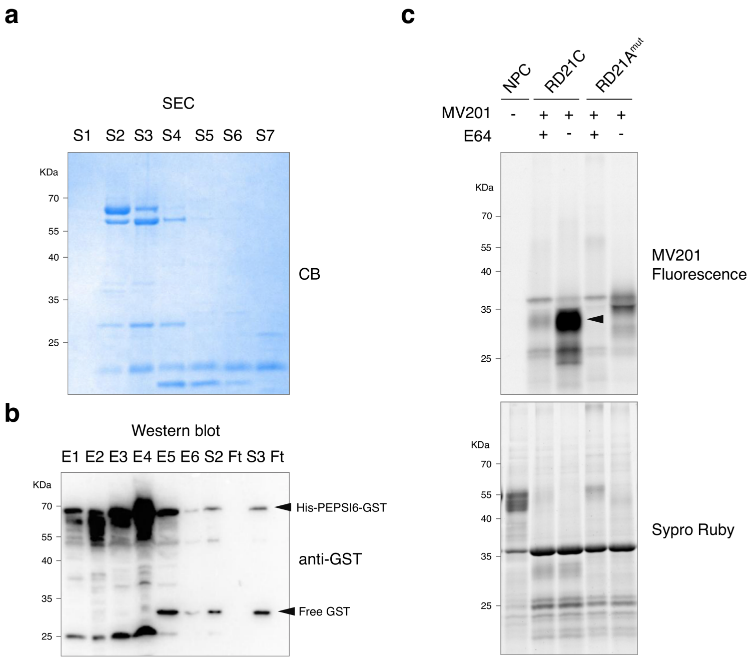


**Extended Data Fig. 11. Production of recombinant His–PEPSI6–GST and validation of RD21C protease activity. a**, SDS–PAGE analysis of recombinant His–PEPSI6–GST following Ni–NTA affinity purification and subsequent size-exclusion chromatography (SEC). Protein purity and integrity were assessed by Coomassie Brilliant Blue (CB) staining. S1-S7 denote individual SEC fractions. **b**, Immunoblot analysis using an anti-GST antibody of fractions collected during protein purification, including Ni–NTA elution fractions (**E1–E6**) and size-exclusion chromatography (SEC) fractions, including flow-through (**FT**) and selected fractions (**S2 and S3**). **c**, Activity-based protein profiling (ABPP) of papain-like cysteine proteases in apoplastic fluid isolated from *N. benthamiana* leaves expressing RD21C or the catalytically inactive RD21A mutant. Active proteases were labeled with the fluorescent activity-based probe MV201 in the presence or absence of the cysteine protease inhibitor E64. Fluorescent signals correspond to probe-labeled active RD21C (black arrowhead), whereas labeling is reduced upon E64 treatment confirming specificity or in samples expressing RD21A mutant (background signals from *N. benthamiana*). Total protein loading is shown by Sypro® Ruby staining.

**Supplementary Video 1.** 4D live imaging of PEPSI6-GFP (green) and the plastid-targeted tdTomato showing colocalization and dynamic behavior over 4 hours, acquired at 15-s intervals for 998 time points (10,978 total frames). Each frame represents a merge of 11 z-planes spanning a total depth of 5 µm, acquired using a piezo stage. Scale bar, 10 µm.
